# StarFunc: fusing template-based and deep learning approaches for accurate protein function prediction

**DOI:** 10.1101/2024.05.15.594113

**Authors:** Chengxin Zhang, Quancheng Liu, Lydia Freddolino

**Author notes:** To whom correspondence should be addressed. Correspondences may also be addressed to Chengxin Zhang. These two authors contributed equally (Chengxin Zhang, Quancheng Liu).

## Abstract

Deep learning has significantly advanced the development of high-performance methods for protein function prediction. Nonetheless, even for state-of-the-art deep learning approaches, template information remains an indispensable component in most cases. While many function prediction methods use templates identified through sequence homology or protein-protein interactions, very few methods detect templates through structural similarity, even though protein structures are the basis of their functions. Here, we describe our development of StarFunc, a composite approach that integrates state-of-the-art deep learning models seamlessly with template information from sequence homology, protein-protein interaction partners, proteins with similar structures, and protein domain families. Large-scale benchmarking and blind testing in the 5^th^ Critical Assessment of Function Annotation (CAFA5) consistently demonstrate StarFunc’s advantage when compared to both state-of-the-art deep learning methods and conventional template-based predictors.

## INTRODUCTION

The functions of proteins arise in large part from each protein’s specific tertiary structure, making structure-based prediction a promising avenue for function prediction. We previously developed COFACTOR (1,2), a function prediction server that incorporates annotations from structural templates identified from TM-align (3), one of the fastest global structure alignment tool at the time. In addition to structure templates, COFACTOR also incorporates function annotations from BLASTp/PSI-BLAST sequence homologs and protein-protein interaction (PPI) partners. COFACTOR participated in the third Critical Assessment of Function Annotation challenge (CAFA3), which aimed to benchmark state-of-the-art algorithms to predict Gene Ontology (GO) terms. In CAFA3, COFACTOR was the first ranked method in the limited knowledge Biological Process category, and also achieved high ranks in several other categories (4).

Despite the promise of structure-based function prediction, aside from a few notable exceptions (1,5,6), the vast majority of existing function prediction methods ignore tertiary structure, regardless of whether they use traditional template-based approaches (7,8) or state-of-the-art deep learning models (9-15). There are two main obstacles to structure-based function prediction. First, until very recently (16,17), consistent and scalable prediction of protein structures at near experimental accuracies had not been possible. This not only causes the lack of high-quality structure models to be unavailable for many target proteins of interest, but also makes many proteins with experimental function annotation inaccessible as templates for structure-based function prediction. Second, performing structural searches through structure template databases were typically very time consuming. For example, even with the relatively fast TM-align engine that performs a pairwise alignment within approximately 0.1 second, it still takes COFACTOR several hours to scan through the BioLiP (18) database of experimental structures with function annotations.

Fortunately, recent advances in structure bioinformatics have largely mitigated both challenges noted above. First, novel deep learning algorithms, especially AlphaFold2 (16) and the associated AlphaFold Protein Structure Database (AFDB) (19), have made accurate structure models available for a huge number of biologically significant proteins. Second, recent innovations in ultrafast and sensitive database search algorithms, especially Foldseek (20), have made structure template identification more than 1000-fold faster than TM-align.

To take advantage of recent progress in structural bioinformatics, we introduce StarFunc, a new composite approach to structure-based protein function prediction. Compared to its predecessor COFACTOR, StarFunc has made several important advancements. First, its structure-based component adds a fast Foldseek-based structure prefiltering stage to select the subset of related templates for full length TM-align alignment, providing both the efficiency of Foldseek and the sensitivity of TM-align for structural template detection. Second, the structure template database is expanded to include not only experimental structures from the BioLiP database (18) but also computational models from AFDB (19), providing a far larger library of possible structural templates. Third, a newly developed deep learning component and another Pfam family-based component method are added to further improve the prediction accuracy. According to the most recent CAFA5 challenge and an independent test, StarFunc has achieved state-of-the-art performance with GO prediction accuracy exceeding many existing template-based and deep learning predictors.

## MATERIAL AND METHODS

The StarFunc pipeline consists of five complementary component methods (**Figure 1A**), each of which can independently predict GO terms for a target protein structure, and which are subsequently combined to provide a consensus annotation set. Below we outline the component pipelines, how they were derived, and the combination rules used to obtain the final StarFunc predictions.

**Figure 1.**
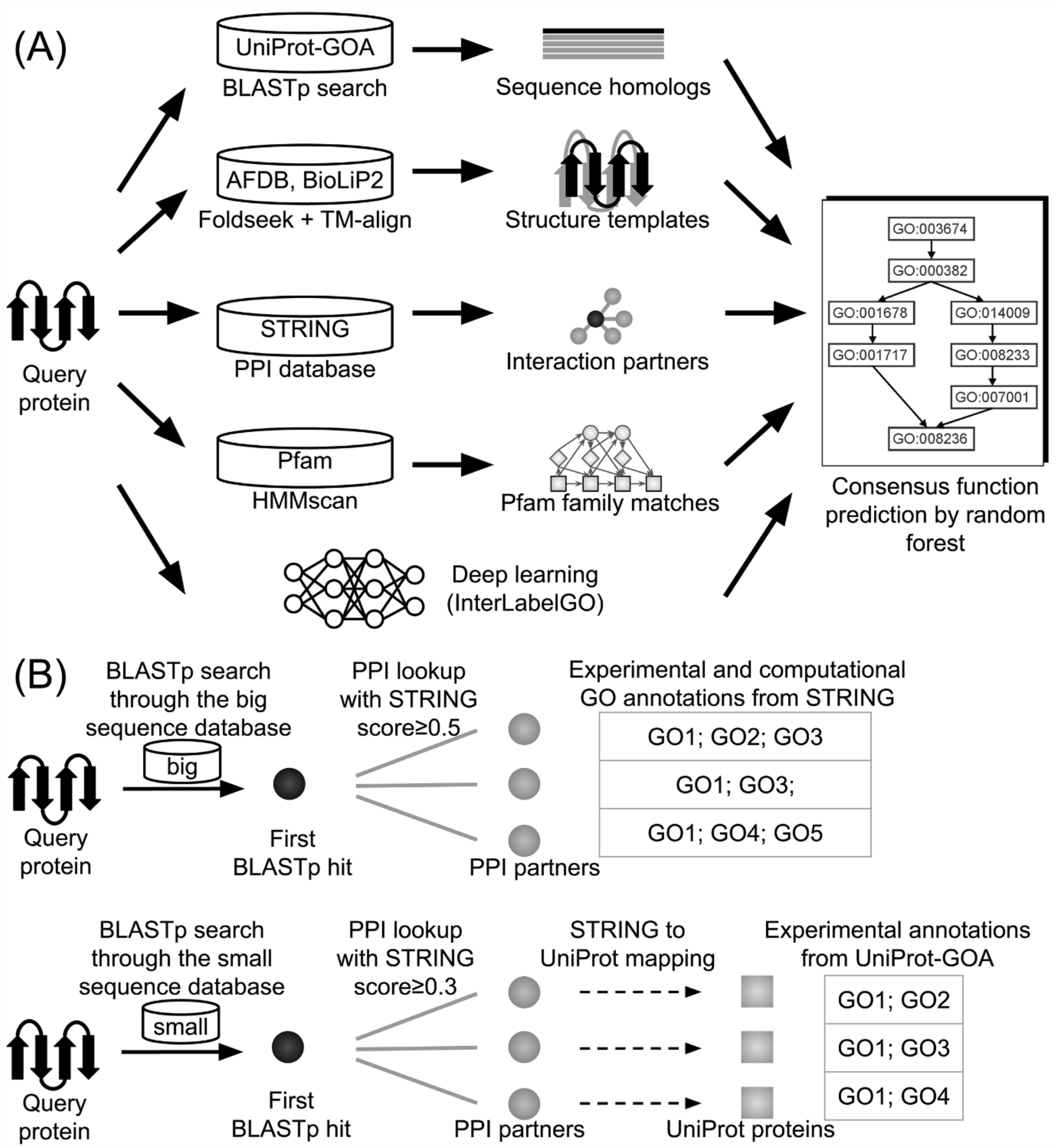
**(A)** Flowchart of StarFunc, which consists of five component methods based on sequence homologs, structure templates, protein-protein interaction partners, Pfam family matches, and deep learning. **(B)** Flowchart of StarFunc’s PPI component, which derives function predictions from both a “big” and a “small” PPI database, as described in the main text.

### Component method 1: sequence-based annotation

Starting from the target protein structure, StarFunc first extracts its sequence, which is searched against the UniProt Gene Ontology Annotation (UniProt-GOA) database by BLASTp to identify homologous templates with function annotations. The prediction score for each GO term *q* is calculated as:

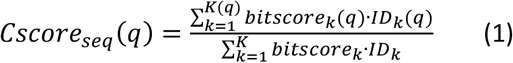

In this equation, *bitscore*_*k*_ and *ID*_*k*_ are the bit-score and sequence identity of the *k*-th BLASTp hit, while *K* is the total number of hits with at least one GO term in the same GO aspect (Molecular Function, MF; Biological Process, BP; or Cellular Component, CC) as term *q*. Meanwhile, *bitscore*_*k*_ (*q*), *ID*_*k*_ (*q*) and *K*(*q*) are the corresponding values for templates with GO term *q*. Here, the sequence identity for hit *k* is calculated as:

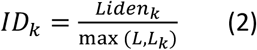

Here, *L* and *L*_*k*_ are the length of the query and template sequence respectively, while *Liden*_*k*_ are the number of identical amino acids between the two proteins in the BLASTp alignment. The design of *Cscore*_*seq*_(*q*) follows our recent benchmark study (21), which indicated that such a scoring function combining both the bit-score and sequence identity outperforms previous scoring functions (1,2,10) that use bit-score or sequence identity only.

When generating the UniProt-GOA sequence database for StarFunc, we only use experimental GO annotations as defined by CAFA (4) (evidence codes: EXP, IDA, IPI, IMP, IGI, IEP, HTP, HDA, HMP, HGI, HEP, TAS, and IC), excluding computational prediction such as those with evidence code IEA. Negative annotations indicating lack of protein function (with qualifier “NOT” or evidence codes “ND”) are also excluded. Parent GO terms are automatically propagated from the leaf terms annotated by UniProt-GOA. Similar to the practice in previous CAFA challenges (4), protein binding-only templates, i.e., template proteins whose only Molecular Function (MF) leaf term is GO:0005515 “protein binding”, are not used for MF term prediction, because “protein binding” is a highly generic function description. Protein binding-only templates are also not used by other component methods in StarFunc.

### Component method 2: structure-based annotation

The query protein structure is searched through two databases: the BioLiP2 database (18) containing functions for chains derived from experimental protein structures, and the subset of computational models deposited in the AFDB (19) that overlaps with experimentally annotated proteins from UniProt-GOA. The initial structural search is performed by Foldseek with an E-value cutoff of 10. This is followed by re-alignment of Foldseek hits using TM-align to obtain full length alignments and the associated TM-scores. The list of structure templates identified in AFDB and BioLiP2 are concatenated together and the prediction score for GO term *q* is obtained via:

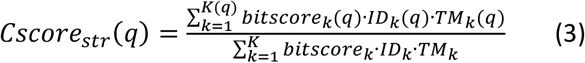

Here, *bitscore*_*k*_ is the bit-score of Foldseek alignment to the *k*-th template; *ID*_*k*_ is the sequence identity calculated from the TM-align alignment. *TM*_*k*_ is the TM-score between query and template structures. If the query and template have different sequence lengths, the TM-score normalized by the longer of the two structures will be used.

### Component method 3: protein-protein interaction (PPI)-based annotation

The third component method derives protein functions from the query protein’s PPI partner as defined by the PPI networks in the STRING database (22). Here, STRING is a comprehensive database of experimentally characterized and computationally predicted pairwise PPIs, where each PPI is represented in the following format:

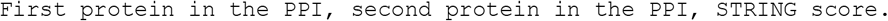

Here, the STRING score is the likelihood of a PPI and ranges from 0 to 1. Since PPIs are symmetric, if STRING has an entry for protein A interacting with protein B, it will also have another entry for protein B interacting with protein A at the same STRING score. Although STRING proteins indeed have GO annotations, the STRING database does not differentiate between experimental annotations and computational annotations. On the other hand, it is possible to map STRING proteins to UniProt accessions, based on which experimental GO annotations can be retrieved from UniProt-GOA. However, only a subset of STRING proteins can be mapped to UniProt unambiguously using the mapping file provided by the STRING database. To address this dilemma, we constructed two PPI databases from the STRING database based on the two different approaches to retrieve GO annotation (**Figure 1B**). A “big” PPI database was constructed by extracting all STRING PPIs with STRING scores ≥0.5 for which the second protein in the PPI has at least one GO term (experimental or computational) annotated by STRING. Another “small” PPI database was constructed by extracting all STRING PPIs with STRING scores ≥ 0.3 for which the second protein can be mapped to a UniProt-GOA protein with experimental annotation. The “big” database is approximately 27 times larger than the “small” database, but the latter has higher quality GO annotations. For both the “big” and “small” databases, the sequence of the first protein in each PPI was collected to make the corresponding sequence databases, for query proteins to be matched against.

During function prediction, the query sequence is searched separately through the “big” and “small” sequence databases. The BLASTp hit with the best bit-score in each sequence database is used to find its PPI partners. The sequence identities between the query sequence and the best hit in the “big” and “small” sequence databases are denoted as *ID*_*big*_ and *ID*_*small*_, respectively. The PPI-derived function prediction for a GO term *q* is then be calculated as:

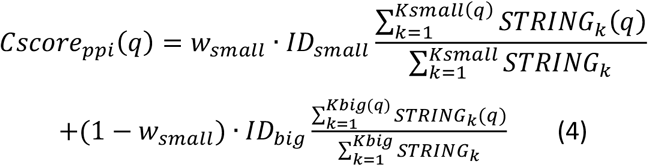

*Ksmall* and *Kbig* are the total number of PPI partners identified in the “small” and “big” PPI databases, respectively. *STRING*_*k*_ is the STRING score for the *k*-th interaction partner. *Ksmall*(*q*), *Kbig*(*q*) and *STRING*_*k*_ (*q*) are the corresponding values of the subset of PPI partners with GO term *q*. Meanwhile, *w*_*ppi*_ is the weight to balance between predictions from the “small” and “big” databases. It is calculated as:

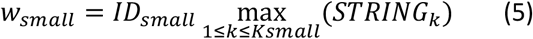

### Component method 4: Pfam-based annotation

In this component method, the query protein is searched through the hidden Markov models (HMMs) from the Pfam database (23) by the HMMsearch tool from the HMMER package (24) using options “-Z 61295632 -E 1000”, which are the HMMER parameters used to construct the Pfam database. The prediction for GO term *q* is obtained by:

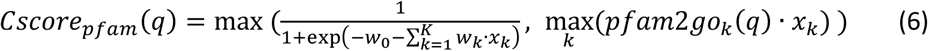

Here, *w*_0_ and *w*_*k*_ are parameters of a GO term-specific logistic regression model trained by mlpack (25) using options “--max_iterations 100 --tolerance 1e-4 --lambda 0.5”. *x*_*k*_ *=* 1 − *E*_*k*_/1000 indicates the match of the query sequence to the *k*-th Pfam family, where *E*_*k*_ is the E-value from HMMsearch. In practice, although the full Pfam database has 19632 families, we only use the most frequent *K*=7000 Pfam families in this component method. The indicator function *pfam*2*go*_*k*_ (*q*) equals 1 or 0 depending on whether the *k*-th Pfam family is associated with GO term *q* in the Pfam2GO mapping file (26) provided by the Gene Ontology Consortium.

### Component method 5: deep learning-based annotation (InterLabelGO)

In addition to the above component methods that rely on template information, StarFunc also incorporates InterLabelGO, a deep learning component for template-free function prediction (**Figure S1**). InterLabelGO is inspired by our earlier deep learning-based GO prediction tool ATGO (10) but with significant improvements in both feature extraction and loss function design. Technical details for training and architecture design of the InterLabelGO model will be described elsewhere (Liu Q, Zhang C, Freddolino PL, manuscript in preparation). Briefly, starting from the query protein sequence, InterLabelGO uses the ESM2 protein language model (27) (version esm2_t36_3B_UR50D) to extract sequence features in the form of an *L*×2560 embedding matrices, where 2560 is the number of embedding dimensions. Similar to ATGO, three different embedding matrices corresponding to the last three layers of the ESM2 model are extracted. Each embedding matrix is average-pooled along the sequence to derive an embedding vector of length 2560. Each of the three embedding vectors is separately fed to a single layer fully connected feedforward neural network with Gaussian Error Linear Unit (GELU) activation function. Each of the three feedforward networks has an output of length 1024. The outputs are concatenated to generate an embedding vector of length 3072. This embedding vector is fed to another feedforward network with two hidden layers to perform multi-class classification. Each output value is transformed by a sigmoid function to scale between 0 and 1. The output contains the prediction scores for the 1571, 4871 and 1066 most frequent GO terms in the MF, BP and CC aspects, respectively. These numbers of GO terms are chosen such that there are at least 50 training proteins for each BP term and at least 10 training proteins for each MF or CC term. In total, 5 different InterLabelGO models are trained for each of three GO aspects, and the prediction scores of the 5 models are averaged.

### StarFunc consensus prediction

The final consensus function predictions of StarFunc are derived using random forest (RF) models. To this end, for each of the three GO aspects (MF, BP and CC), we trained five different models using five-fold cross-validation. Each model consists of 4000 trees and was trained by the LightGBM package using a learning rate of 0.002. For each GO term *q*, the input features include the five prediction scores for term *q* from all five component methods, as well as the “naïve” probability of term *q*, i.e., the number of UniProt-GOA proteins with GO term *q* divided by the number of all UniProt-GOA proteins with experimental annotations in the same GO aspect. These six features are used by each RF model to perform a binary classification. A consensus prediction score is derived from the average RF output scores from the five RF models.

Although the non-machine learning-based component methods (sequence-, structure- and PPI-based methods) always follow the true path rule, where the prediction score of a child term never exceeds its parent terms, this is not guaranteed for machine learning components (Pfam- and deep learning-based component methods as well as the final RF models). To address this issue, after deriving the consensus predictions, we propagate the prediction scores from the child terms towards the root term such that a parent term will have a prediction score that is no less than any of its child terms.

In practice, applying all RF models to classify all >43000 possible GO terms would be very time consuming. Therefore, for each of the three GO aspects, the final RF-based consensus prediction only considers the subset of GO terms that are among the top 100 terms with the highest prediction scores in at least one of the five component methods.

### Performance evaluation

Following conventions used in the Critical Assessment of Function Annotation (CAFA) challenges (4), we evaluated the performance of protein function predictions using three metrics: the maximum F-measure (Fmax), the maximum of information accretion-weighted F-measure (wFmax), and the minimal semantic distance (Smin). Fmax is the primary evaluation metric for CAFA challenges round 1, 2 and 3 (4), and is defined as:

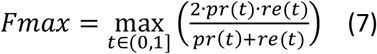

Here, *pr*(*t*) and *re*(*t*) are the precision and recall, respectively, at the prediction score threshold t. They are defined as:

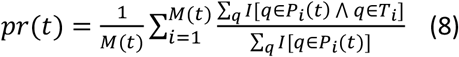

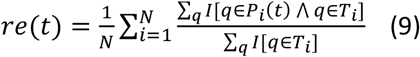

Here, *N* is the total number of proteins in the test set; *M*(*t*) is the number of proteins with at least one predicted GO term with prediction score ≥ t. *T*_*i*_ is the set of experimentally determined (ground truth) GO terms, including their parent terms, for protein *i. P*_*i*_ (*t*) is the set of predicted GO terms for protein i with prediction score ≥ t. *I*[] is the indicator function (i.e., the Iverson’s bracket). For both *T*_*i*_ and *P*_*i*_ (*t*), the root terms of the three GO aspects (GO:0003674 “molecular_function”, GO:0008150 “bio-logical_process”, and GO:0005575 “cellular_component”) are excluded. Additionally, for MF prediction, targets for which the only ground truth MF leaf term is GO:0005515 “protein binding” are excluded from MF evaluation, as this term is very common and, in isolation, not highly informative (28).

An alternative scoring measure is the wFmax, the main evaluation metric for CAFA5. The wFmax is defined as:

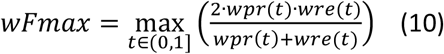

Here, *wpr*(*t*) and *wre*(*t*) are the precision and recall, respectively, weighted by the information accretions (29) of GO terms. They are defined as:

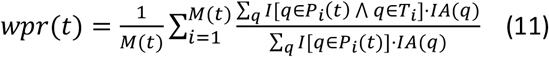

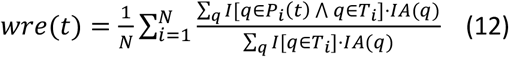

The information accretion, also known as the information content, for GO term *q* is defined as:

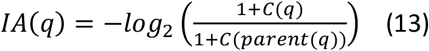

Here, *C*(*q*) is the number of proteins with GO terms *q* among experimentally annotated proteins in the UniProt-GOA database; *C*(*parent*(*q*)) is the number of UniProt-GOA database proteins with all parent terms of GO term *q*.

A related information accretion-weighted metric is Smin, which is calculated as:

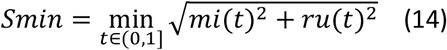

Here, *mi*(*t*) and *ru*(*t*) are the misinformation and remaining uncertainty at prediction score threshold *t*. They are defined as:

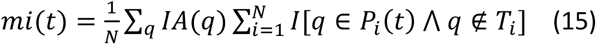

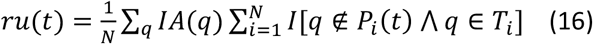

While both wFmax and Fmax range between 0 and 1, with a higher value indicating higher accuracy, Smin ranges between 0 and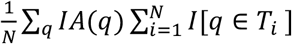], with a lower value indicating more accurate prediction.

In CAFA1 to CAFA3, the evaluation metrics (Fmax, wFmax, and Smin) for the three GO aspects (MF, BP and CC) were considered separately. In the most recent CAFA5 challenge, the average wFmax over the three GO aspects was used as the primary scoring criterion. Therefore, for each scoring metric we report both the GO aspect-specific results and the average metric values over the three GO aspects.

## RESULTS

### Dataset

We trained and tested StarFunc using the same Gene Ontology definition as in the recent CAFA5 challenge (i.e., the “go-basic.obo” file released on 2023-01-1). Following CAFA5, the training dataset uses UniProt-GOA release 2022-11-17. The 142,311 experimental annotated proteins from this UniProt-GOA release are used to generate the template databases for StarFunc’s sequence- and structure-based component method, the “small” sequence database for the PPI-based component method, and the training dataset for the Pfam- and deep learning-based component methods. The template database also used PPI information from STRING database version 11.5 and Pfam version 35.0, both versions being the latest version of the respective databases prior to the 2022-11-17 time cutoff.

To generate a validation dataset, we collected all 2041 proteins with new annotations in UniProt-GOA release 2023-03-16 but no annotation in the same GO aspect as of UniProt-GOA 2022-11-17. This validation dataset was used to tune the hyperparameters of the component methods. Additionally, GO predictions from all component methods on the validation dataset were used to train StarFunc’s final consensus RF models by five-fold cross-validation.

We also generated an independent test set by collecting all 2475 proteins with new annotations in UniProt-GOA release 2023-05-18 but no annotation in the same GO aspect as of UniProt-GOA 2023-03-16. This is the dataset used to compare the performance of StarFunc and other existing function predictors.

### Overall performance

To assess the predictive performance of StarFunc, we compared its predictive accuracy on our independent test set to three traditional template-based methods (COFACTOR (1), GoFDR (7), and GOtcha (8)) and eight state-of-the-art deep learning methods (SPROF-GO (9), ATGO+ (10), TALE+ (11), DeepGOplus (12), AnnoPRO (14), ProteInfer (15), DeepGO-SE (13) and DeepFRI (5)). For all three GO aspects, StarFunc consistently and significantly outperforms the other existing methods (two-tail paired t-test p-value<0.01; see **Figure 2**) with 12.1%, 8.3% and 8.8% higher wFmax for MF, BP and CC, respectively. Similar conclusions are reached based on the Fmax and Smin metrics (**Table S1**).

**Figure 2.**
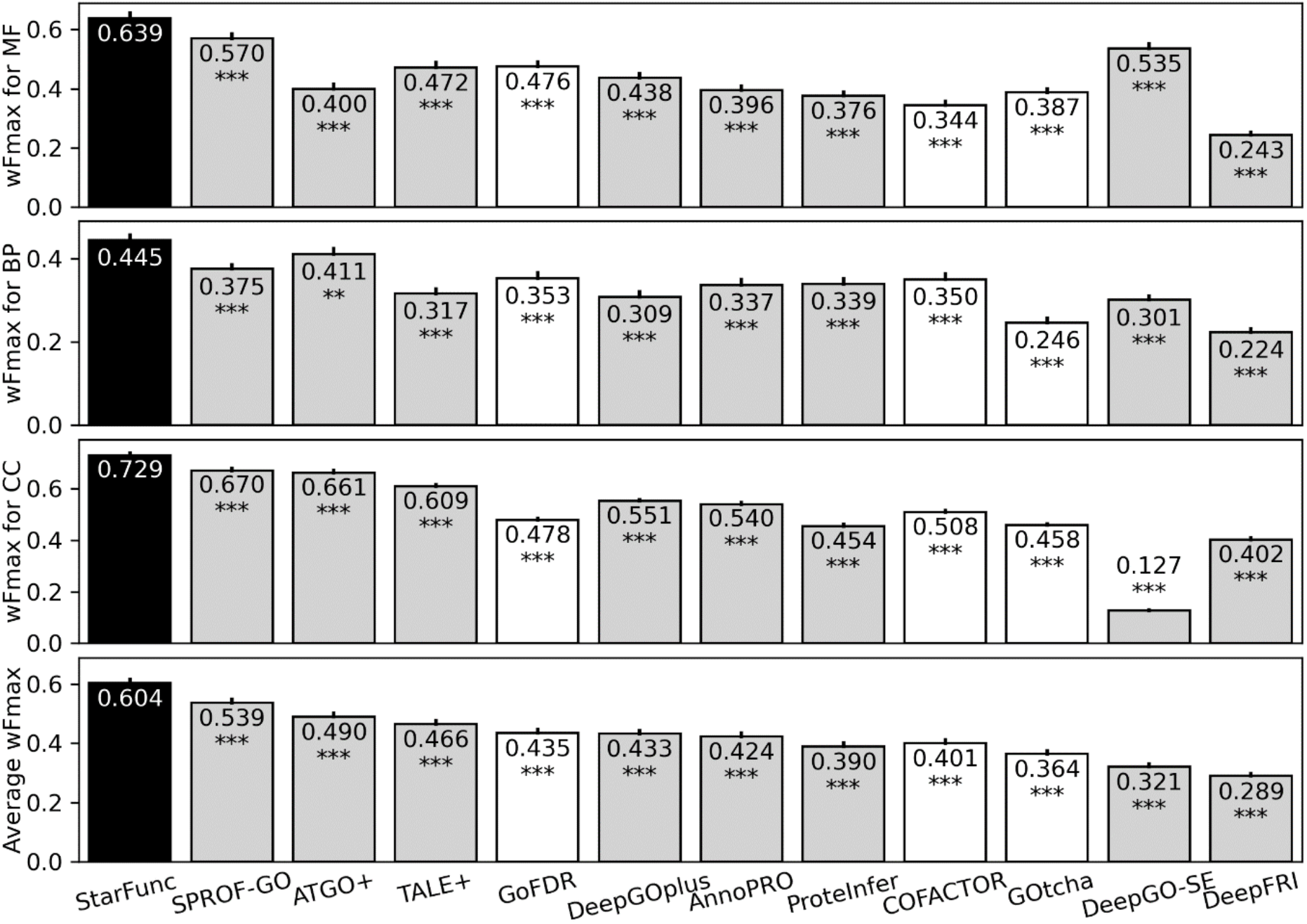
Overall performance of StarFunc compared with existing programs, as measured by the wFmax of the three aspects of GO and the average wFmax over the three aspects. Methods are ranked in descending order of the average wFmax. White and grey colors indicate existing programs that are non-deep learning and deep learning-based, respectively. Error bars at the top of each bar plot indicate the standard error of mean (SEM) for wFmax values. The ** and *** signs indicate p-0.001<p-value≤0.01, and p-value≤0.001, respectively. To calculate the p-values, we calculate the wFmax values of each protein target for each method; these per-target values are used to calculate p-values by a two-tailed paired t-test between StarFunc and each comparison program.

It is noteworthy that although both StarFunc and COFACTOR perform structure alignment through large structure template databases, their speed is very different. For a medium size protein of 400 residues, the average running time for COFACTOR and StarFunc were approximately 10 hours (2) and 16 minutes (**Figure S2**), respectively. The high speed of StarFunc arises largely from the Foldseek prefiltering in the structure-based component, as expensive full length structure alignment by TM-align only needs to be perform for a small subset (usually <100) template structures that with significant Foldseek alignments; this approach allows StarFunc to run much more quickly despite searching a 8-fold larger database of protein structures.

A perhaps surprising observation from our testing is that although the four most accurate algorithms all use deep learning (StarFunc, SPROF-GO, ATGO+ and TALE+), template-based methods (GoFDR, COFACTOR, and GOtcha) can still rival or outperform some deep learning methods (DeepGO-SE and DeepFRI). One notable difference between the four most accurate deep learning methods (StarFunc, SPROF-GO, ATGO+ and TALE+) and the two least accurate deep learning methods (DeepGO-SE and DeepFRI) is that the former set uses sequence homology search as a component, while the latter set are fully template-free pure machine learning methods. These discrepancies in overall performance suggested that template information is still an indispensable component of protein function prediction even for deep learning methods.

Among all tested deep learning methods, DeepFRI has the lowest average wFmax despite being the only deep learning model that incorporates tertiary structure information. This is probably because DeepFRI relies on relatively conventional convolutional network architecture and did not use more advanced transformer-based models as in other methods (9-13).

### CAFA performance

A preliminary version of StarFunc participated in the most recent CAFA5 challenge that officially closed in December 2023 under the group name hfm7zc. Since the InterLabelGO pipeline was still underdevelopment at the time of CAFA5 (and, indeed, a development version of it participated independently in CAFA5 as group “Evans”), the CAFA5 version of StarFunc used SPROF-GO instead of InterLabelGO as its deep learning component, resulting in an approximately 3.6% lower wFmax values than the final version of CAFA5 (**Table S1**). Despite being not fully developed at the time, StarFunc still manage to rank 5^th^ among 1625 CAFA5 teams from 96 countries based on the average wFmax (https://www.kaggle.com/competitions/cafa-5-protein-function-prediction/leaderboard).

### Ablation study

To understand factors contributing to the high accuracy of StarFunc, we checked the performance of each individual component and the effect of their removal on final model performance. Individually speaking, the most accurate component method is InterLabelGO, followed in order by the sequence-based, structure-based, Pfam-based, and PPI-based components (**Figure 3A**). It is notable that the consensus prediction shows significantly higher performance than any individual component, even the deep learning based InterLabelGO pipeline. However, the fact that the PPI-based component has the worst accuracy does not mean it is the least useful component. In fact, in an ablation study where each component was removed from StarFunc, the most significant drop of performance was caused by exclusion of InterLabelGO and the PPI-based component (**Figure 3B**). On the other hand, removal of the sequence-based component method, despite being the second most accurate component, has only a minimal effect on the final accuracy. This is probably due to the redundancy of the sequence-based component to other components such as the structure-based component method, whereas the PPI pipeline provides more unique information to inform the consensus.

**Figure 3.**
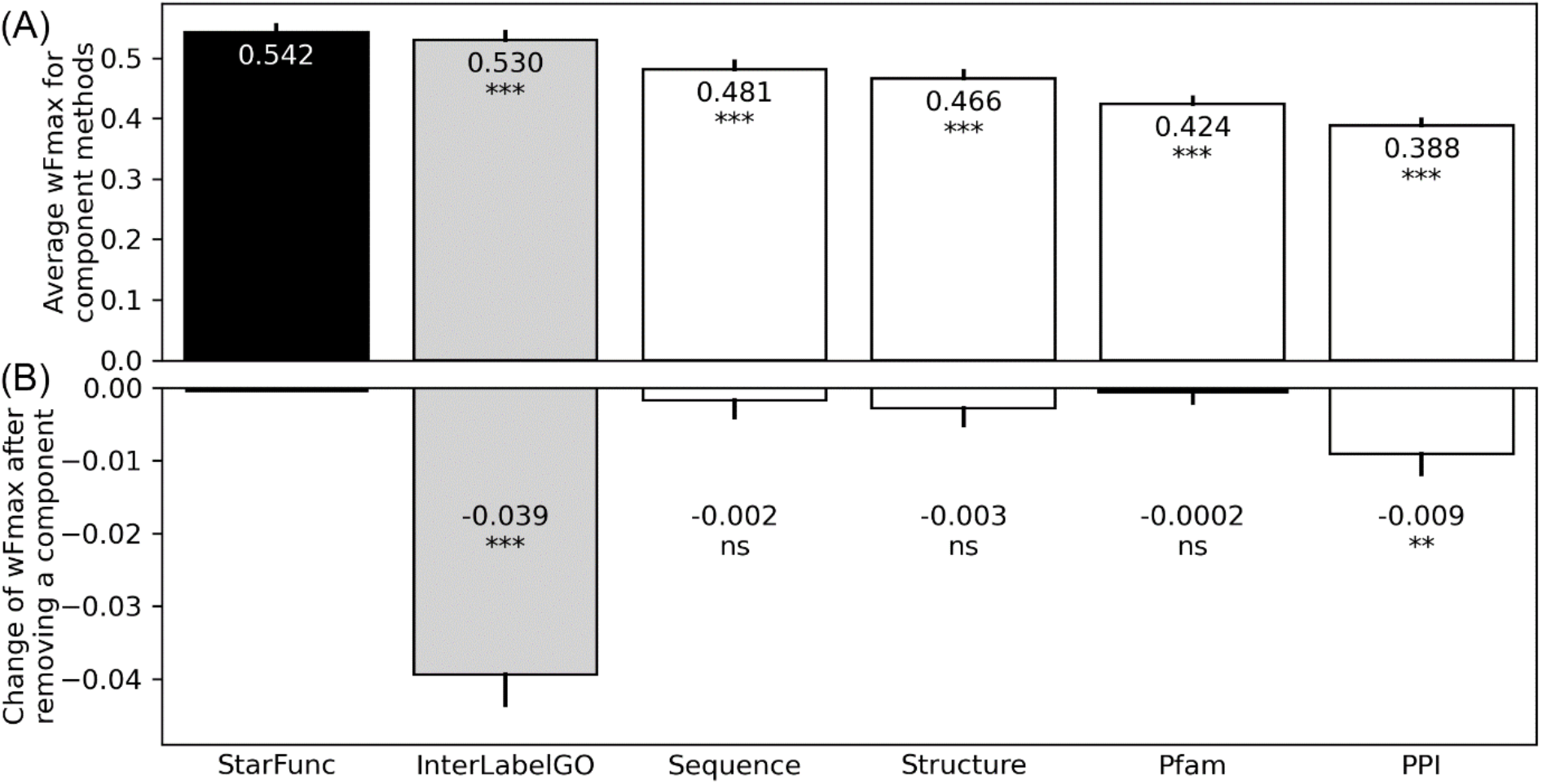
Ablation study to quantify the importance of different component methods of StarFunc. **(A)** Average wFmax over the three GO aspects for each of the individual component methods and the consensus prediction. **(B)** Decrease of average wFmax after removing each component method. White and grey colors indicate component methods that are non-deep learning and deep learning-based, respectively. Error bars at the top of each bar plot indicate the standard error of mean (SEM) for wFmax values. The ns, ** and *** signs indicate p-value>0.05, 0.001<p-value≤0.01, and p-value≤0.001, respectively, where p-values are calculated by two-tailed paired t-test between the consensus StarFunc pipeline and a component method (considering the weighted F-measure values for each protein as the individual data points).

### WEB SERVER

#### Server input

The mandatory input for the StarFunc web server at https://seq2fun.dcmb.med.umich.edu/StarFunc/ is a PDB-format structure file for a single-chain protein structure. Selenomethionine (MSE) in the PDB file is converted to methionine (MET) while heteroatoms of other non-standard amino acids and small molecules are ignored. If multiple chains or multiple models are present in the input file, only the first chain of the first model will be considered. The sequence of the target protein is converted from the “ATOM” records of the PDB file rather than extracting from its “SEQRES” record. The user is also highly encouraged to provide their email addresses so that they will be notified upon job completion; otherwise, they need to bookmark a randomly generated link for server output.

#### Server output

The output webpage contains six annotation panels (**Figure 4**). The first panel shows the sequence and structure of the input protein (**Figure 4A**), while the remaining panels show template information and predicted protein functions.

**Figure 4.**
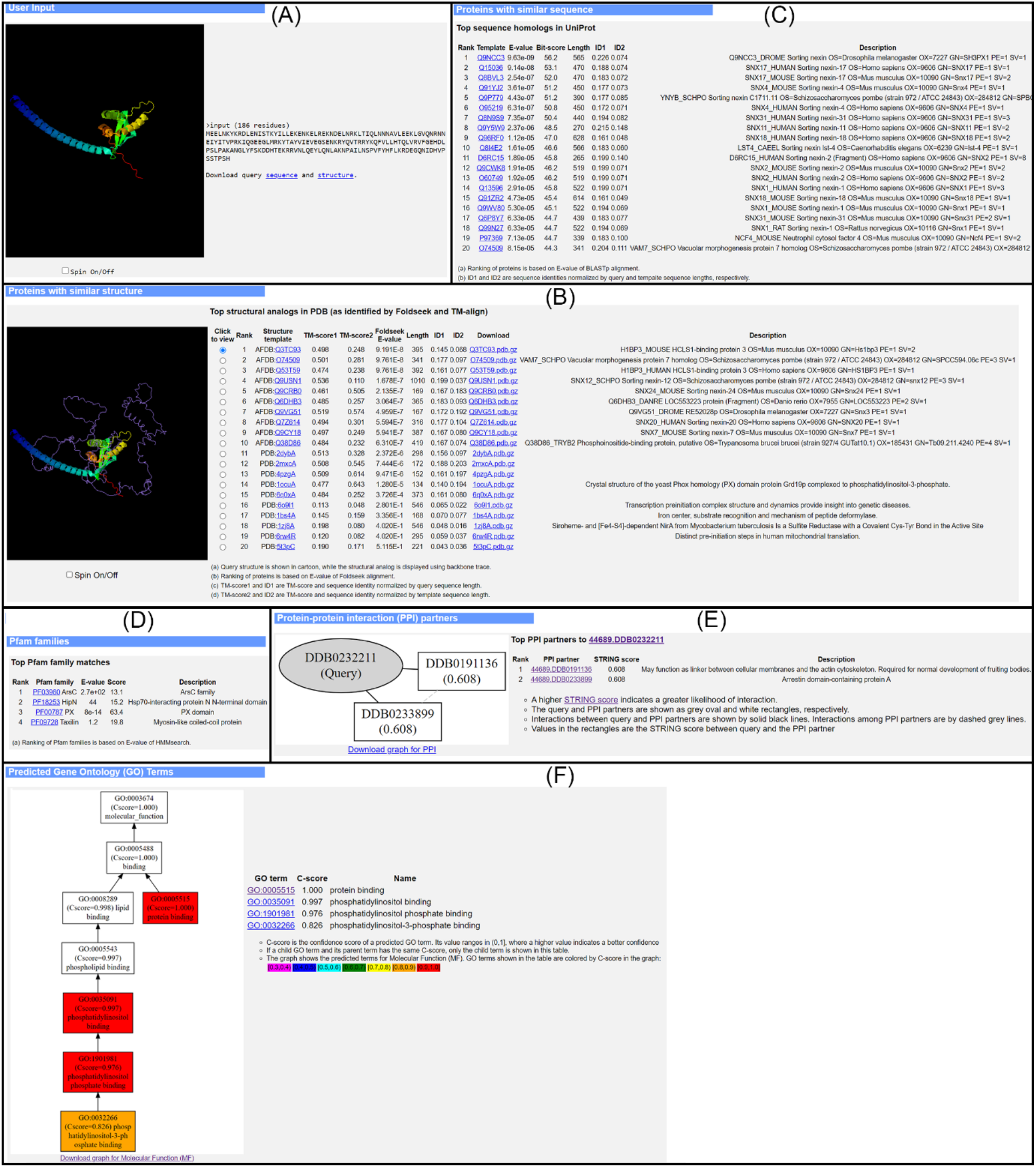
Example output of StarFunc for the SnxA protein from *Dictyostelium discoideum* (UniProt accession Q54GY1). **(A)** Input sequence and structure. **(B)** Structure templates. **(C)** Sequence templates. **(D)** Pfam families. **(E)** PPI partners. **(F)** Predicted GO terms. While the actual web page shows three separate panels for MF, BP and CC, this figure only shows the MF panel due to space limitations.

The second panel lists the top 20 structure templates from AFDB and BioLiP2, together with their TM-scores, Foldseek E-values, sequence identities calculated from the TM-align alignment, and a description of the protein drawn from either the UniProt database (for AFDB templates) or the PDB database (for BioLiP2 templates) (**Figure 4B**). The template structure (purple ribbon) superimposed onto the query structure (rainbow cartoon) is shown in a JSmol applet (30) on the left hand side of the panel.

The third panel lists the top 20 hits from a BLASTp search through UniProt-GOA, together with their E-values, bit-scores, sequence identities and description from UniProt (**Figure 4C**).

The fourth panel shows the matched Pfam families as well as the scores and E-values reported by HMMsearch (**Figure 4D**).

The fifth panel shows all PPI partners in the STRING database with STRING score≥0.5, along with their STRING scores and descriptions. An interaction graph showing the PPIs among the target protein and the PPI partners is drawn using Graphviz (31) at the left-hand side of the panel (**Figure 4E**).

The last panel shows the predicted GO terms in three separate tables which correspond to the MF, BP, and CC aspects of GO (**Figure 4F**). Within each table, the GO terms, as well as their predicted scores and descriptions, are sorted in descending order of predicted scores. If both a parent term and a child term have the same prediction score, only the child term is listed. Meanwhile, a directed acyclic graph for the predictions of each GO aspect is drawn using Graphviz at the left-hand side.

As an illustrative example, we show the StarFunc prediction of the SnxA protein (UniProt accession: Q54GY1, **Figure 4**) from our test set. SnxA is a highly selective phosphatidylinositol-3,5-bisphosphate binding protein (GO:0080025 “phosphatidylinositol-3,5-bisphosphate binding”) from the phagocytic vesicle membrane of the social amoeba *Dictyostelium discoideum* (32). Although StarFunc is not able to predict this highly specific term, it is able to predict its parent term GO:1901981 “phosphatidylinositol phosphate binding” with a very high prediction score of 0.976 (**Figure 4F**). However, the prediction scores for GO:1901981 is quite modest by most component methods (0.496, 0.350, 0.262, and 0.167 by InterLabelGO, structure, sequence and Pfam, respectively) except for the PPI-based component, which predicts the term with an appreciable score of 0.620. This is due to the detection of PPI partner Arrestin domain-containing protein A (STRING ID: 44689.DDB0233899, **Figure 4E**), which binds to phosphatidylinositol phosphate. This example shows that even the PPI-based method, which has the worst performance on its own among all StarFunc components, can still make an important contribution to accurate final prediction; it also provides an excellent example of the consensus prediction being far stronger than that of any individual component.

It should be noted that the RF consensus models used in the final stage of StarFunc do not merely aggregate predictions from individual components. Instead, these models will distinguish correct predictions from mispredictions when performing the consensus. For example, the FhuE protein (UniProt accession: P16869) is a transporter of iron across the outer membrane of *E. coli* (**Figure 5A**) (33). The StarFunc component methods predict both the correct BP term GO:1901678 “iron coordination entity transport” (**Figure 5B**) and the incorrect BP term GO:0043604 “amide biosynthetic process” (**Figure 5C**). For both terms, the highest prediction scores come from InterLabelGO, which predicts GO:1901678 and GO:0043604 with scores of 0.807 and 0.669, respectively. The sequence-based component also predicts both terms with scores of 0.376 and 0.485, respectively. This is because the top sequence hit protein, *Pseudomonas aeruginosa* FpvA (UniProt accession: P48632) is a bifunctional protein where the N-terminal domain is involved in the biosynthesis of pyoverdine, an amide-containing compound (GO:0043604), while the rest is a transporter of iron (GO:1901678) (34). In the final StarFunc prediction, the score for the correct term GO:1901678 is increased to 0.921 while that for the incorrect term GO:0043604 decreased to 0.468. This causes the final StarFunc prediction to be much more accurate than any of its component methods (**Figure 5D**).

**Figure 5.**
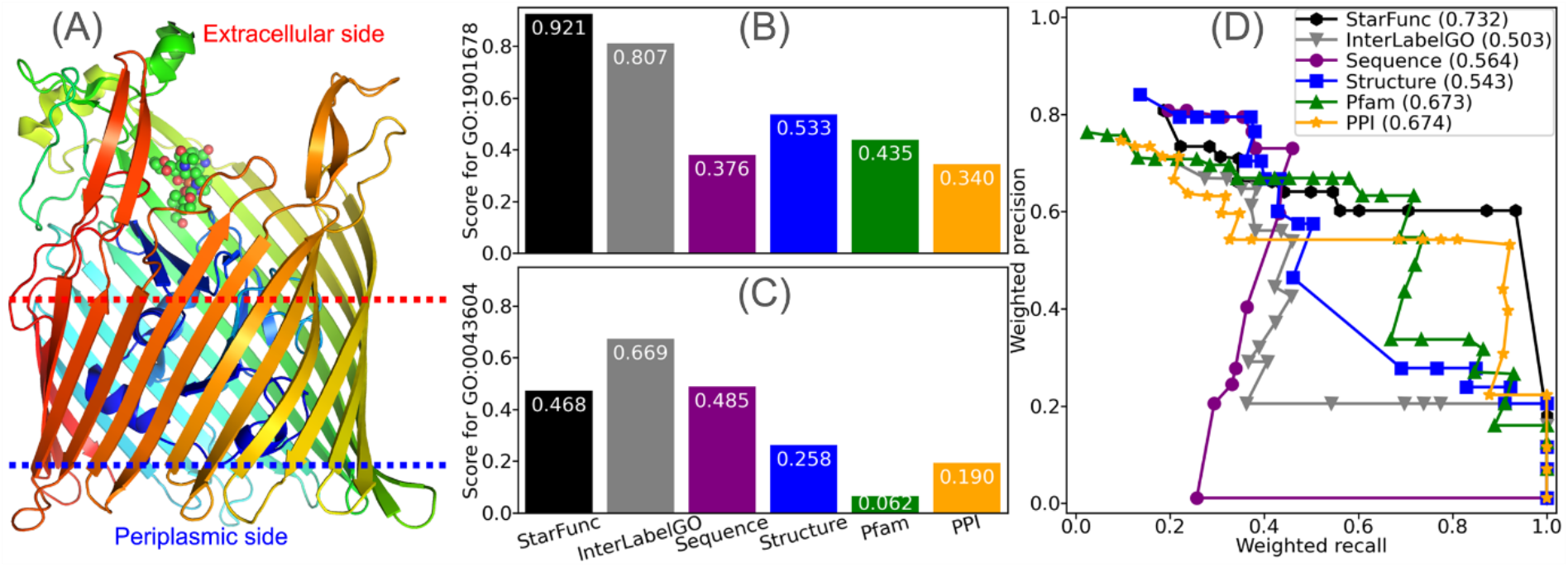
StarFunc prediction for *Escherichia coli* FhuE. **(A)** Structure of FhuE (PDB ID: 6e4v) shown in rainbow, where blue to red indicates N-to C-terminus. Spheres show coprogen, which is a small molecule that binds iron and is transported by FhuE. **(B-C)** Prediction scores for **(B)** GO:1901678 “iron coordination entity transport” and **(C)** GO:0043604 “amide biosynthetic process” by StarFunc and its component methods. **(D)** Information accretion weighted precision and recall by StarFunc and its component methods. Values in the parentheses are the wFmax values.

### Function prediction for the human proteome reveals putative ubiquitination-related proteins

To apply StarFunc to the biomedically-important topic of detecting unknown biological roles of human proteins, we applied StarFunc to all 20389 proteins from the human reference proteome curated in the neXtProt (35) database release 2023-09-11. The full set of predictions are available at https://seq2fun.dcmb.med.umich.edu/StarFunc/HUMAN. Among these 20389 proteins, there are 18397 Protein Evidence 1 (PE1) proteins whose existence is unequivocally confirmed experimentally, e.g., by mass spectroscopy or fluorescence immunostaining. Another 1381 proteins are “missing proteins” whose existences are either confirmed at transcript level (PE2) or predicted computationally (PE3 and PE4). The remaining 611 proteins are PE5 proteins, i.e., uncertain or dubious sequences such as pseudogenes or erroneous translation products.

To study what kinds of functions are more common among missing proteins (PE2, PE3 and PE4) than PE1 proteins, we applied a rate-ratio test (36) for GO terms predicted by StarFunc (**Figure 6A**) or annotated by UniProt-GOA (**Figure 6B**). Based on MF and BP annotations from both StarFunc and UniProt-GOA, among missing proteins, the most prominent function category is olfactory receptors (GO:0004984 “olfactory receptor activity”) which are usually GPCRs (GO:0004930 “G protein-coupled receptor activity”) responsible for smell and taste. Around one third of missing proteins falls within this category, which is consistent with previous reports (37). This is not surprising given that these GPCRs are embedded in the membrane and not susceptible to trypsin digestion and subsequent mass spectroscopy-based protein detection; it is also likely that many odorant-specific GPCRs are expressed in only small numbers of cells.

**Figure 6.**
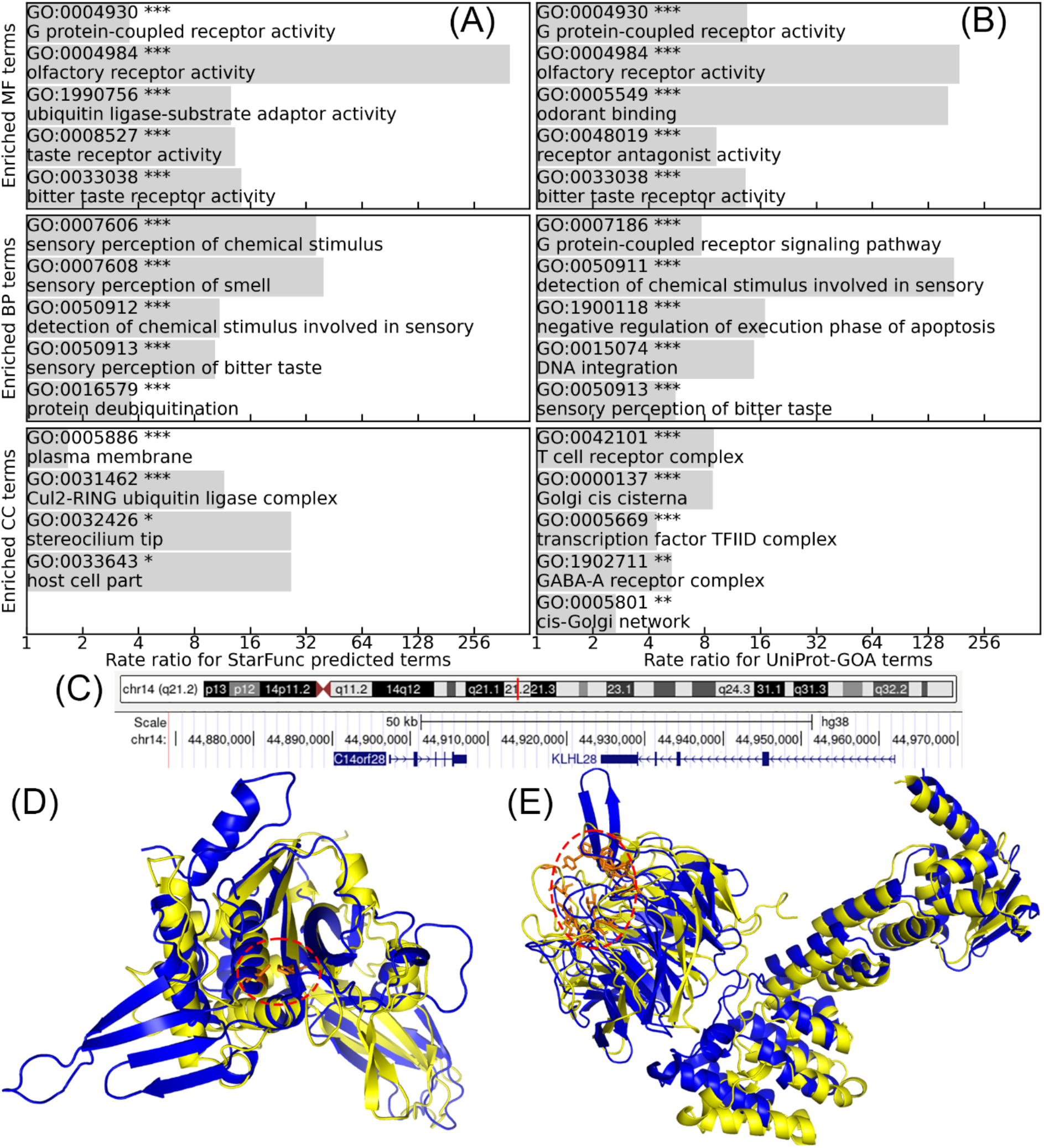
Identification of annotations that are enriched in missing proteins relative to PE1 proteins according to a rate ratio test. **(A)** StarFunc predicted GO terms with prediction score≥0.9. **(B)** UniProt-GOA annotated GO terms, including computational GO terms with IEA evidence. Only the five most significantly enriched terms are shown. The *, ** and *** signs indicate 0.01<p-value≤0.05, 0.001<p-value≤0.01, and p-value≤0.001, respectively. **(C)** Gene organization of C14orf28 and KLHL28 on human chromosome 14 shown by USCS genome browser (https://genome.ucsc.edu/cgi-bin/hgTracks?db=hg38&position=chr14%3A44885680%2D44975526) (40). **(D)** Structure superimposition between C14orf28 (blue) and the catalytic domain of UBP8 (yellow, residue 145 to 445), whose catalytic residues (residues 154 and 395) are shown in orange sticks within the red dashed circle. **(E)** Structure superimposition between KLHL28 (blue) and KLHL20 (yellow), whose substrate binding residues (residues 326, 329, 356, 373, 378, 405, 421, 423, 451, 498, 499, 515, 545, 567, 592) are shown in orange sticks within the red dashed circle.

On the other hand, the enriched CC terms among StarFunc predictions are quite distinct from those found in UniProt-GOA annotations. In particular, there are 12 missing proteins with similar sequences from the “Preferentially expressed antigen of melanoma” (PRAME) family (**Figure S3** and **Table S3**) that are annotated as GO:0031462 “Cul2-RING ubiquitin ligase complex”, which is an important complex targeting proteins for ubiquitination-dependent protein degradation. This prediction is mainly caused by a PE1 protein from the PRAME family, PRAME family member 9 (UniProt accession: P0DUQ2), which has GO:00031462 annotation and high identity (>90%) to these 12 missing proteins.

In addition to missing proteins, there are also four PE1 proteins with unknown functions (uPE1 proteins) that are predicted by both the template-based and deep learning components of StarFunc to associate with ubiquitination (**Table 1**). The first two proteins, RNF222 (UniProt accession: A6NCQ9) and RNF224 (UniProt accession: P0DH78) are both predicted to have GO:0004842 “ubiquitin-protein transferase activity”. For both proteins, all top templates from the sequence and structure components have low sequence identities (<30%) and structural similarities (TM-score<0.5) but almost all of them are ubiquitin-protein ligases (**Figure S4** and **Figure S5**). Since the prediction scores from the sequence- and structure-based methods are based on consensus of multiple templates, the prediction scores from both component methods are high (>0.5). The other two proteins are C14orf28 and KLHL28 (UniProt accession: Q4W4Y0 and Q9NXS3, respectively). Their genes are located adjacent to each other on chromosome 14 (**Figure 6C**), indicating a likely functional relation. C14orf28 and KLHL28 have significant structural similarity to UBP8, a deubiquitinase (UniProt accession: Q09738, Figure 6D) and KLHL20, a component of an E3 ubiquitin ligase complex (UniProt accession: Q9Y2M5, Figure 6E), respectively. Therefore, C14orf28 and KLHL28 are predicted by StarFunc to have GO:0019783 “ubiquitin-like protein peptidase activity” and GO:0016567 “protein ubiquitination” with prediction scores of 0.931 and 0.941, respectively. These examples demonstrate how StarFunc enables the detection of likely additional human proteins involved in an important and widely studied process (i.e., protein ubiquitination), serving as a launch point for additional biological studies.

**Table 1.** Summary of uPE1 human proteins predicted to be associate with ubiquitination.

| UniProt<br>accession | Protein<br>name | Gene<br>name | GO term | Prediction scores |  |  |  |  |  |
| --- | --- | --- | --- | --- | --- | --- | --- | --- | --- |
|  |  |  |  | Star-Func | Struc-<br>ture | Seq-<br>uence | PPI | Pfam | Inter-<br>labelGO |
| A6NCQ9 | RING finger<br>protein 222 | RNF222 | GO:0004842<br>“ubiquitin-protein<br>transferase activity” | 1.000 | 0.593 | 0.541 | 0.000 | 0.780 | 0.952 |
| P0DH78 | RING finger<br>protein 224 | RNF224 | GO:0004842<br>“ubiquitin-protein<br>transferase activity” | 1.000 | 0.602 | 0.574 | 0.000 | 0.048 | 0.941 |
| Q4W4Y0 | C14orf28 | C14orf28 | GO:0019783<br>“ubiquitin-like protein<br>peptidase activity” | 0.931 | 0.824 | 0.000 | 0.000 | 0.000 | 0.635 |
| Q9NXS3 | Kelch-like<br>protein 28 | KLHL28 | GO:0016567 “protein<br>ubiquitination” | 0.941 | 0.598 | 0.497 | 0.243 | 0.376 | 0.961 |

## DISCUSSION AND CONCLUSION

In this study, we developed StarFunc, a composite approach to protein function prediction that combines annotations from sequence homologs, structure templates, protein interactions, Pfam families, and deep learning in a unified framework. The final consensus prediction of StarFunc consistently outperforms all of its individual component methods as well as other state-of-the-art algorithms in the field. Its execution speed far exceeds its predecessor, COFACTOR, that also uses structure template alignment. Detailed ablation study highlights the significant contributions from the newly added deep learning and the redesigned protein interaction-based components.

One limitation of StarFunc is the lack of a text mining component, which is shown by recent studies to significantly boost annotation accuracies when literatures are available (38). Another shortcoming is the limited types of features in the deep learning component, which can potentially be boosted by features extracted from sequence alignments (39) and tertiary structure (5). Future versions of StarFunc will try to improve the performance following these two directions.

## SUPPLEMENTARY DATA

Supplementary Data are available at NAR online.

### ACKNOWLEDGEMENT

The authors thank Dr Xiaoqiong Wei for insightful discussions. This work used the Advanced Cyberinfrastructure Coordination Ecosystem: Services & Support (ACCESS) program, which is supported by National Science Foundation [2138259, 2138286, 2138307, 2137603, and 2138296].

## FUNDING

This work was supported by the National Institute of Allergy and Infectious Diseases [AI134678 to P.L.F.]. The funders had no role in study design, data collection and analysis, decision to publish, or preparation of the manuscript. Funding for open access charge: National Institutes of Health.

## CONFLICT OF INTEREST

None declared.

## Notes

### Competing Interest Statement

The authors have declared no competing interest.

