## Supplemental Figures and Tables for "StarFunc: fusing template-based and deep learning approaches for accurate protein function prediction"

#### Table of Contents

**Supporting Figures**


**Figure S1.** Architecture of InterLabelGO.
**Figure S2.** Running time of the full StarFunc pipeline.
**Figure S3.** Multiple sequence alignment of PRAME family proteins among "missing" human proteins that are predicted to be associated with the Cul2-RING ubiquitin ligase complex.
**Figure S4.** Structure and sequence templates used for StarFunc prediction of A6NCQ9.
**Figure S5.** Structure and sequence templates used for StarFunc prediction of P0DH78.
**Figure S6.** Structure and sequence templates used for StarFunc prediction of Q4W4Y0.
**Figure S7.** Structure and sequence templates used for StarFunc prediction of Q9NXS3.


**Supporting Tables**


**Table S1.** Fmax, Smin and wFmax values for the test set.
**Table S2.** Fmax, Smin and wFmax values for StarFunc and its component methods on the validation set.
**Table S3.** Summary of PRAME family proteins among missing proteins that are predicted by StarFunc to be associated with the Cul2-RING ubiquitin ligase complex.

### Supporting Figures

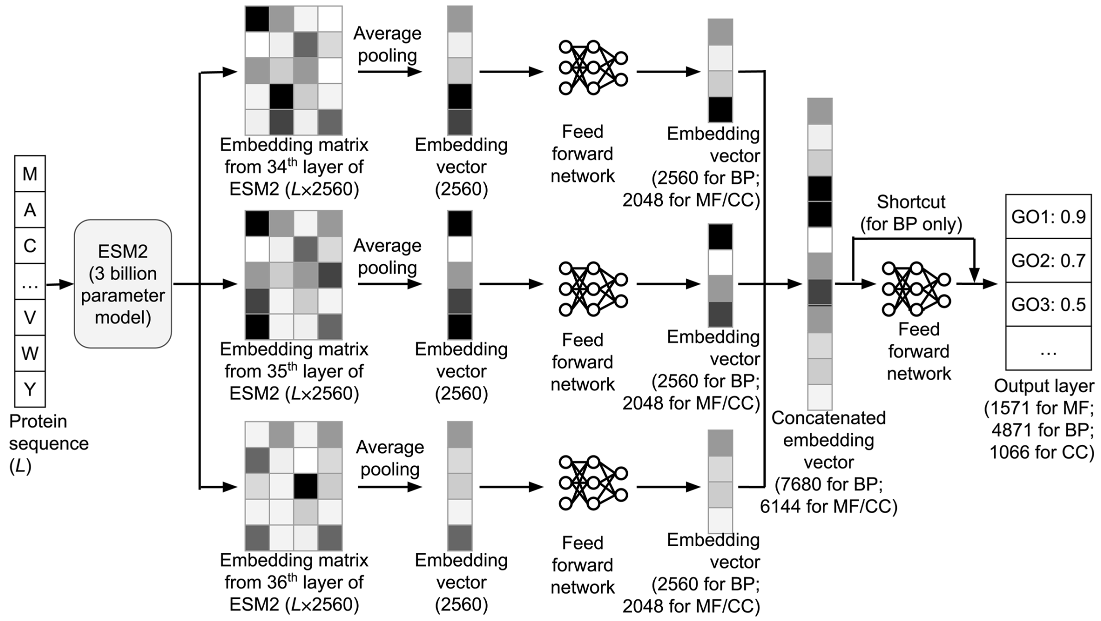


**Figure S1.** Architecture of InterlabelGO, the deep learning-based function prediction component of StarFunc. The dimensions of different layers are shown in the parenthesis. Each InterlabelGO model is specific for one of the three GO aspects (MF, BP and CC). The hyperparameters of models for different GO aspects, especially the dimensions of the feed forward networks, are different. These differences are caused by the much larger number of BP terms, which require more complex models than MF and CC.

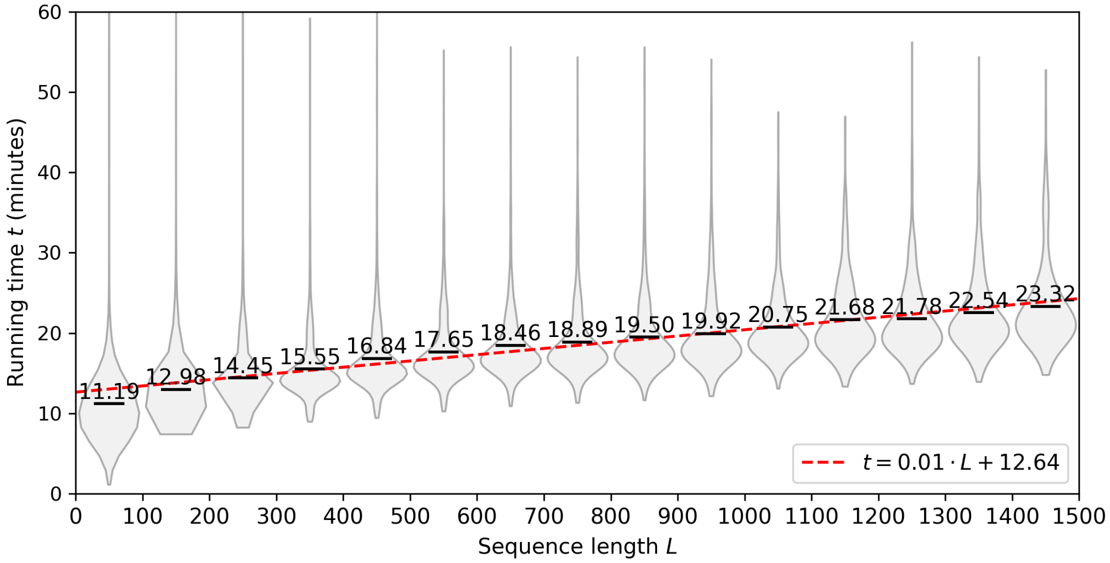


**Figure S2.** Running time of the full StarFunc pipeline for all 19598 human proteins with up to 1500 residues from the AlphaFold database, stratified by sequence length into 100 amino acid bins. Black horizontal lines for each violin indicate the mean running time for each protein sequence length bin. The red dashed line is the least square fit for running time versus protein sequence length.

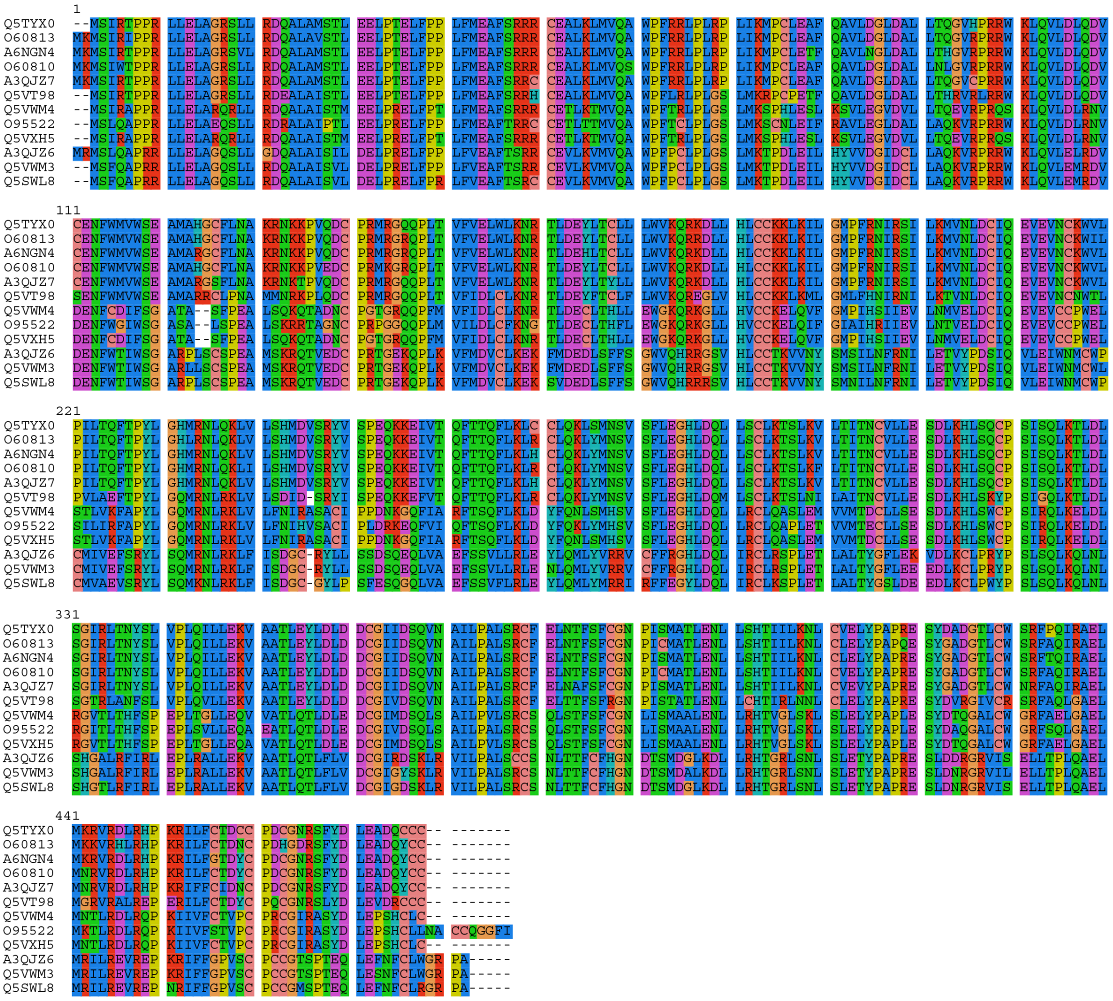

**Figure S3.** Multiple sequence alignment of PRAME family proteins among "missing" human proteins that are predicted to be associated with the Cul2-RING ubiquitin ligase complex.

#### Proteins with similar structure

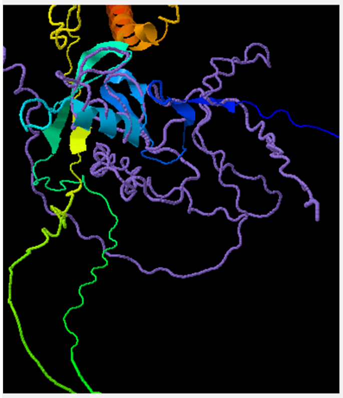

Spin On/Off

##### Top structural analogs (as identified by Foldseek and TM-align)

| Click to view | Rank | Structure template | TM-score1 | TM-score2 | Foldseek E-value | Length | ID1 | ID2 | Download | Description |
|---|---|---|---|---|---|---|---|---|---|---|
| | 1 | AFDB:O64824 | 0.201 | 0.195 | 3.361E-3 | 227 | 0.186 | 0.181 | O64824.pdb.gz | O64824_ARATH E3 ubiquitin-protein ligase RMA OS=Arabidopsis thaliana OX=3702 GN=At2g23780 PE=1 SV=1 |
| | 2 | AFDB:Q9Y7K6 | 0.203 | 0.065 | 1.020E-1 | 741 | 0.150 | 0.045 | Q9Y7K6.pdb.gz | YGI4_SCHPO Uncharacterized RING finger protein C2A9.04c OS=Schizosaccharomyces pombe (strain 972 / ATCC 24843) OX=284812 GN=SPBC2A9.04c PE=4 SV=1 |
| | 3 | AFDB:Q96A61 | 0.164 | 0.122 | 1.512E-1 | 297 | 0.123 | 0.091 | Q96A61.pdb.gz | TRI52_HUMAN E3 ubiquitin-protein ligase TRIM52 OS=Homo sapiens OX=9606 GN=TRIM52 PE=1 SV=1 |
| | 4 | AFDB:Q9HEV3 | 0.147 | 0.115 | 1.724E-1 | 320 | 0.205 | 0.141 | Q9HEV3.pdb.gz | Q9HEV3_EMEND GATA factor AREB beta OS=Emericella nidulans OX=162425 GN=areB PE=4 SV=1 |
| | 5 | PDB:5dkaA | 0.176 | 0.388 | 1.964E-1 | 96 | 0.073 | 0.167 | 5dkaA.pdb.gz | A C2HC zinc finger is essential for the RING-E2 interaction of the ubiquitin ligase RNF125. |
| | 6 | PDB:5a31B | 0.172 | 0.422 | 2.391E-1 | 84 | 0.077 | 0.202 | 5a31B.pdb.gz | Atomic Structure of the Apc/C and its Mechanism of Protein Ubiquitination. |
| | 7 | PDB:5l9uB | 0.169 | 0.483 | 2.912E-1 | 71 | 0.068 | 0.211 | 5l9uB.pdb.gz | |
| | 8 | PDB:7b5lH | 0.193 | 0.104 | 3.109E-1 | 423 | 0.068 | 0.035 | 7b5lH.pdb.gz | Ubiquitin ligation to F-box protein targets by SCF-RBR E3-E3 super-assembly. |
| | 9 | PDB:5gm6a | 0.166 | 0.287 | 5.994E-1 | 123 | 0.059 | 0.106 | 5gm6a.pdb.gz | Structure of a yeast activated spliceosome at 3.5 angstrom resolution |
| | 10 | AFDB:Q9LX93 | 0.203 | 0.153 | 7.305E-1 | 301 | 0.168 | 0.123 | Q9LX93.pdb.gz | ATL55_ARATH E3 ubiquitin-protein ligase RING1 OS=Arabidopsis thaliana OX=3702 GN=ATL55 PE=1 SV=1 |
| | 11 | AFDB:Q68DV7 | 0.180 | 0.056 | 8.330E-1 | 783 | 0.223 | 0.063 | Q68DV7.pdb.gz | RNF43_HUMAN E3 ubiquitin-protein ligase RNF43 OS=Homo sapiens OX=9606 GN=RNF43 PE=1 SV=1 |
| | 12 | AFDB:Q9M371 | 0.137 | 0.048 | 9.498E-1 | 790 | 0.159 | 0.044 | Q9M371.pdb.gz | Q9M371_ARATH Extra-large G-like protein, putative (DUF3133) OS=Arabidopsis thaliana OX=3702 GN=F15G16.60 PE=1 SV=1 |
| | 13 | AFDB:M9PE32 | 0.148 | 0.029 | 1.014 | 1314 | 0.168 | 0.028 | M9PE32.pdb.gz | M9PE32_DROME Fife, isoform D OS=Drosophila melanogaster OX=7227 GN=Fife PE=1 SV=1 |
| | 14 | AFDB:Q5A432 | 0.147 | 0.053 | 1.083 | 688 | 0.118 | 0.038 | Q5A432.pdb.gz | GAT1_CANAL Transcriptional regulatory protein GAT1 OS=Candida albicans (strain SC5314 / ATCC MYA-2876) OX=237561 GN=GAT1 PE=2 SV=1 |
| | 15 | AFDB:Q5CCK4 | 0.162 | 0.066 | 1.083 | 780 | 0.155 | 0.044 | Q5CCK4.pdb.gz | VAL2_ARATH B3 domain-containing transcription repressor VAL2 OS=Arabidopsis thaliana OX=3702 GN=VAL2 PE=1 SV=1 |
| | 16 | PDB:7oikA | 0.167 | 0.009 | 1.234 | 4426 | 0.064 | 0.003 | 7oikA.pdb.gz | E3 ubiquitin ligase RNF213 employs a non-canonical zinc finger active site and is allosterically regulated by ATP |
| | 17 | PDB:2lgvA | 0.162 | 0.332 | 1.502 | 100 | 0.086 | 0.190 | 2lgvA.pdb.gz | Selective recruitment of an e2~ubiquitin complex by an e3 ubiquitin ligase. |
| | 18 | PDB:7ns4b | 0.137 | 0.090 | 1.604 | 340 | 0.041 | 0.026 | 7ns4b.pdb.gz | GID E3 ligase supramolecular chelate assembly configures multipronged ubiquitin targeting of an oligomeric metabolic enzyme. |
| | 19 | PDB:6ft6u | 0.141 | 0.195 | 1.713 | 150 | 0.041 | 0.060 | 6ft6u.pdb.gz | Structure of the nuclear exosome captured on a maturing preribosome. |
| | 20 | PDB:8a3tU | 0.167 | 0.307 | 1.953 | 114 | 0.059 | 0.114 | 8a3tU.pdb.gz | Cryo-EM structure of the S. cerevisiae APC/C-Cdh1 complex |

(a) The query structure is shown in cartoon, while the structural analog is displayed using a backbone trace.
(b) Ranking of proteins is based on E-value of Foldseek alignment.
(c) TM-score1 and ID1 are TM-score and sequence identity normalized by query sequence length.
(d) TM-score2 and ID2 are TM-score and sequence identity normalized by template sequence length.

#### Proteins with similar sequence

##### Top sequence homologs in UniProt

| Rank | Template | E-value | Bit-score | Length | ID1 | ID2 | Description |
|---|---|---|---|---|---|---|---|
| 1 | Q9FY48 | 8.08e-06 | 48.5 | 1625 | 0.109 | 0.015 | KEG_ARATH E3 ubiquitin-protein ligase KEG OS=Arabidopsis thaliana OX=3702 GN=KEG PE=1 SV=2 |
| 2 | Q96D59 | 1.02e-05 | 47.0 | 192 | 0.114 | 0.130 | RN183_HUMAN E3 ubiquitin-protein ligase RNF183 OS=Homo sapiens OX=9606 GN=RNF183 PE=1 SV=2 |
| 3 | Q8QZS5 | 1.26e-05 | 46.6 | 190 | 0.118 | 0.137 | RN183_MOUSE E3 ubiquitin-protein ligase RNF183 OS=Mus musculus OX=10090 GN=Rnf183 PE=1 SV=1 |
| 4 | F4HY36 | 1.43e-05 | 47.8 | 1805 | 0.132 | 0.016 | F4HY36_ARATH Protein translocase subunit SecA OS=Arabidopsis thaliana OX=3702 GN=SECA2 PE=1 SV=1 |
| 5 | Q557I6 | 8.77e-05 | 45.1 | 552 | 0.109 | 0.043 | Q557I6_DICDI RING-type E3 ubiquitin transferase OS=Dictyostelium discoideum OX=44689 GN=mip1-2 PE=4 SV=1 |
| 6 | A0A384LEC9 | 1.24e-04 | 44.3 | 263 | 0.109 | 0.091 | A0A384LEC9_ARATH RING-type domain-containing protein OS=Arabidopsis thaliana OX=3702 GN=At3g29270 PE=1 SV=1 |
| 7 | Q84JZ7 | 1.24e-04 | 44.3 | 263 | 0.109 | 0.091 | Q84JZ7_ARATH At3g29270 OS=Arabidopsis thaliana OX=3702 GN=At3g29270 PE=1 SV=1 |
| 8 | A0A0R4IL45 | 1.27e-04 | 44.7 | 663 | 0.105 | 0.035 | A0A0R4IL45_DANRE RING-type E3 ubiquitin transferase OS=Danio rerio OX=7955 GN=trim32 PE=3 SV=1 |
| 9 | A8WGA9 | 1.37e-04 | 44.7 | 663 | 0.105 | 0.035 | A8WGA9_DANRE RING-type E3 ubiquitin transferase OS=Danio rerio OX=7955 GN=trim32 PE=2 SV=1 |
| 10 | Q498M5 | 1.40e-04 | 44.7 | 735 | 0.100 | 0.030 | SH3R2_RAT E3 ubiquitin-protein ligase SH3RF2 OS=Rattus norvegicus OX=10116 GN=Sh3rf2 PE=1 SV=1 |
| 11 | Q38BI4 | 2.40e-04 | 43.5 | 387 | 0.095 | 0.054 | Q38BI4_TRYB2 RING-type E3 ubiquitin transferase OS=Trypanosoma brucei brucei (strain 927/4 GUTat10.1) OX=185431 GN=Tb10.70.3100 PE=4 SV=1 |
| 12 | Q9C040 | 2.62e-04 | 43.9 | 744 | 0.086 | 0.026 | TRIM2_HUMAN Tripartite motif-containing protein 2 OS=Homo sapiens OX=9606 GN=TRIM2 PE=1 SV=1 |
| 13 | Q9ESN6 | 2.80e-04 | 43.5 | 744 | 0.086 | 0.026 | TRIM2_MOUSE Tripartite motif-containing protein 2 OS=Mus musculus OX=10090 GN=Trim2 PE=1 SV=1 |
| 14 | D3ZQG6 | 3.01e-04 | 43.5 | 744 | 0.086 | 0.026 | TRIM2_RAT Tripartite motif-containing protein 2 OS=Rattus norvegicus OX=10116 GN=Trim2 PE=1 SV=2 |
| 15 | Q13049 | 3.09e-04 | 43.5 | 653 | 0.095 | 0.032 | TRI32_HUMAN E3 ubiquitin-protein ligase TRIM32 OS=Homo sapiens OX=9606 GN=TRIM32 PE=1 SV=2 |
| 16 | Q8CH72 | 4.04e-04 | 43.1 | 655 | 0.095 | 0.032 | TRI32_MOUSE E3 ubiquitin-protein ligase TRIM32 OS=Mus musculus OX=10090 GN=Trim32 PE=1 SV=2 |
| 17 | Q9P1Y6 | 4.32e-04 | 43.1 | 1649 | 0.118 | 0.016 | PHRF1_HUMAN PHD and RING finger domain-containing protein 1 OS=Homo sapiens OX=9606 GN=PHRF1 PE=1 SV=3 |
| 18 | Q8N6D2 | 5.29e-04 | 42.4 | 247 | 0.100 | 0.089 | RN182_HUMAN E3 ubiquitin-protein ligase RNF182 OS=Homo sapiens OX=9606 GN=RNF182 PE=1 SV=1 |
| 19 | Q8TEC5 | 7.07e-04 | 42.4 | 729 | 0.095 | 0.029 | SH3R2_HUMAN E3 ubiquitin-protein ligase SH3RF2 OS=Homo sapiens OX=9606 GN=SH3RF2 PE=1 SV=3 |
| 20 | Q9N3T6 | 7.13e-04 | 42.4 | 468 | 0.123 | 0.058 | Q9N3T6_CAEEL RING-type domain-containing protein OS=Caenorhabditis elegans OX=6239 GN=CELE_Y47G6A.14 PE=1 SV=2 |

(a) Ranking of proteins is based on the E-value of a BLASTp alignment.
(b) ID1 and ID2 are sequence identities normalized by the query and template sequence lengths, respectively.

**Figure S4.** Structure and sequence templates used for StarFunc prediction of A6NCQ9.

#### Proteins with similar structure

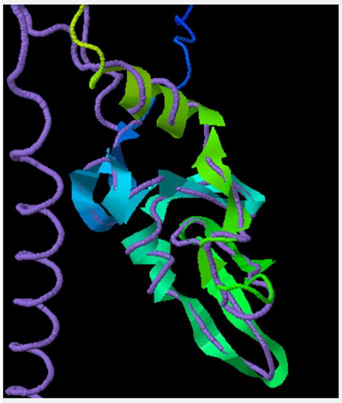

Spin On/Off

**Top structural analogs (as identified by Foldseek and TM-align)**

| Click to view | Rank | Structure template | TM-score1 | TM-score2 | Foldseek E-value | Length | ID1 | ID2 | Download | Description |
|---|---|---|---|---|---|---|---|---|---|---|
| | 1 | AFDB:Q9NXI6 | 0.455 | 0.321 | 3.183E-8 | 227 | 0.212 | 0.145 | Q9NXI6.pdb.gz | RN186_HUMAN E3 ubiquitin-protein ligase RNF186 OS=Homo sapiens OX=9606 GN=RNF186 PE=1 SV=1 |
| | 2 | AFDB:P36406 | 0.458 | 0.131 | 4.595E-7 | 574 | 0.212 | 0.057 | P36406.pdb.gz | TRI23_HUMAN E3 ubiquitin-protein ligase TRIM23 OS=Homo sapiens OX=9606 GN=TRIM23 PE=1 SV=1 |
| | 3 | AFDB:Q8N6D2 | 0.476 | 0.310 | 1.143E-6 | 247 | 0.231 | 0.146 | Q8N6D2.pdb.gz | RN182_HUMAN E3 ubiquitin-protein ligase RNF182 OS=Homo sapiens OX=9606 GN=RNF182 PE=1 SV=1 |
| | 4 | AFDB:O01481 | 0.407 | 0.233 | 2.192E-6 | 283 | 0.179 | 0.099 | O01481.pdb.gz | O01481_CAEEL RING-type domain-containing protein OS=Caenorhabditis elegans OX=6239 GN=C06A5.8 PE=1 SV=1 |
| | 5 | AFDB:A0A384LEC9 | 0.425 | 0.262 | 4.204E-6 | 263 | 0.205 | 0.122 | A0A384LEC9.pdb.gz | A0A384LEC9_ARATH RING-type domain-containing protein OS=Arabidopsis thaliana OX=3702 GN=At3g29270 PE=1 SV=1 |
| | 6 | AFDB:E7F1U3 | 0.421 | 0.101 | 8.604E-6 | 774 | 0.224 | 0.045 | E7F1U3.pdb.gz | E7F1U3_DANRE RING-type E3 ubiquitin transferase OS=Danio rerio OX=7955 GN=trim3a PE=1 SV=1 |
| | 7 | AFDB:Q8BG47 | 0.423 | 0.331 | 4.677E-5 | 203 | 0.224 | 0.172 | Q8BG47.pdb.gz | RN152_MOUSE E3 ubiquitin-protein ligase RNF152 OS=Mus musculus OX=10090 GN=Rnf152 PE=1 SV=1 |
| | 8 | AFDB:Q9H8W5 | 0.418 | 0.124 | 4.991E-5 | 580 | 0.199 | 0.053 | Q9H8W5.pdb.gz | TRI45_HUMAN E3 ubiquitin-protein ligase TRIM45 OS=Homo sapiens OX=9606 GN=TRIM45 PE=1 SV=2 |
| | 9 | AFDB:Q2Q1W2 | 0.324 | 0.066 | 9.571E-5 | 868 | 0.199 | 0.036 | Q2Q1W2.pdb.gz | LIN41_HUMAN E3 ubiquitin-protein ligase TRIM71 OS=Homo sapiens OX=9606 GN=TRIM71 PE=1 SV=1 |
| | 10 | AFDB:Q8QZS5 | 0.439 | 0.364 | 9.571E-5 | 190 | 0.224 | 0.184 | Q8QZS5.pdb.gz | RN183_MOUSE E3 ubiquitin-protein ligase RNF183 OS=Mus musculus OX=10090 GN=Rnf183 PE=1 SV=1 |
| | 11 | PDB:8a3tU | 0.264 | 0.340 | 1.102E-2 | 114 | 0.160 | 0.219 | 8a3tU.pdb.gz | Cryo-EM structure of the S. cerevisiae APC/C-Cdh1 complex |
| | 12 | PDB:5dkaA | 0.233 | 0.361 | 3.559E-2 | 96 | 0.135 | 0.219 | 5dkaA.pdb.gz | A C2HC zinc finger is essential for the RING-E2 interaction of the ubiquitin ligase RNF125. |
| | 13 | PDB:4kblA | 0.272 | 0.123 | 4.928E-2 | 395 | 0.128 | 0.051 | 4kblA.pdb.gz | Structure of HHARI, a RING-IBR-RING Ubiquitin Ligase: Autoinhibition of an Ariadne-Family E3 and Insights into Ligation Mechanism. |
| | 14 | PDB:7kzpL | 0.255 | 0.120 | 1.309E-1 | 370 | 0.109 | 0.046 | 7kzpL.pdb.gz | Structure of the FA core ubiquitin ligase closing the ID clamp on DNA. |
| | 15 | PDB:7oniH | 0.257 | 0.119 | 1.491E-1 | 380 | 0.096 | 0.039 | 7oniH.pdb.gz | CUL5-ARIH2 E3-E3 ubiquitin ligase structure reveals cullin-specific NEDD8 activation. |
| | 16 | PDB:7qcdC | 0.281 | 0.178 | 1.491E-1 | 267 | 0.090 | 0.052 | 7qcdC.pdb.gz | Cryo-EM structure of the Smc5/6 holo-complex. |
| | 17 | PDB:7b5lH | 0.170 | 0.067 | 3.051E-1 | 423 | 0.090 | 0.033 | 7b5lH.pdb.gz | Ubiquitin ligation to F-box protein targets by SCF-RBR E3-E3 super-assembly. |
| | 18 | PDB:7od1A | 0.269 | 0.110 | 3.256E-1 | 430 | 0.090 | 0.033 | 7od1A.pdb.gz | CUL5-ARIH2 E3-E3 ubiquitin ligase structure reveals cullin-specific NEDD8 activation. |
| | 19 | PDB:7oikA | 0.326 | 0.014 | 6.664E-1 | 4426 | 0.096 | 0.003 | 7oikA.pdb.gz | E3 ubiquitin ligase RNF213 employs a non-canonical zinc finger active site and is allosterically regulated by ATP |
| | 20 | PDB:3m62A | 0.328 | 0.061 | 1.051 | 955 | 0.045 | 0.007 | 3m62A.pdb.gz | |

(a) The query structure is shown in cartoon, while the structural analog is displayed using a backbone trace.
(b) Ranking of proteins is based on E-value of Foldseek alignment.
(c) TM-score1 and ID1 are TM-score and sequence identity normalized by query sequence length.
(d) TM-score2 and ID2 are TM-score and sequence identity normalized by template sequence length.

#### Proteins with similar sequence

**Top sequence homologs in UniProt**

| Rank | Template | E-value | Bit-score | Length | ID1 | ID2 | Description |
|---|---|---|---|---|---|---|---|
| 1 | O01481 | 4.68e-08 | 52.8 | 283 | 0.167 | 0.092 | O01481_CAEEL RING-type domain-containing protein OS=Caenorhabditis elegans OX=6239 GN=C06A5.8 PE=1 SV=1 |
| 2 | Q96D59 | 6.60e-08 | 51.6 | 192 | 0.244 | 0.198 | RN183_HUMAN E3 ubiquitin-protein ligase RNF183 OS=Homo sapiens OX=9606 GN=RNF183 PE=1 SV=2 |
| 3 | Q9H0X6 | 3.29e-07 | 50.4 | 261 | 0.231 | 0.138 | RN208_HUMAN RING finger protein 208 OS=Homo sapiens OX=9606 GN=RNF208 PE=1 SV=2 |
| 4 | Q8QZS5 | 3.92e-07 | 49.7 | 190 | 0.237 | 0.195 | RN183_MOUSE E3 ubiquitin-protein ligase RNF183 OS=Mus musculus OX=10090 GN=Rnf183 PE=1 SV=1 |
| 5 | F4HY36 | 4.89e-07 | 50.4 | 1805 | 0.186 | 0.016 | F4HY36_ARATH Protein translocase subunit SecA OS=Arabidopsis thaliana OX=3702 GN=SECA2 PE=1 SV=1 |
| 6 | E7F1U3 | 6.96e-06 | 47.0 | 774 | 0.154 | 0.031 | E7F1U3_DANRE RING-type E3 ubiquitin transferase OS=Danio rerio OX=7955 GN=trim3a PE=1 SV=1 |
| 7 | Q58ER5 | 4.22e-05 | 44.7 | 744 | 0.147 | 0.031 | Q58ER5_DANRE RING-type E3 ubiquitin transferase OS=Danio rerio OX=7955 GN=zgc:113099 PE=2 SV=1 |
| 8 | O01482 | 1.26e-04 | 43.1 | 381 | 0.135 | 0.055 | O01482_CAEEL RING-type domain-containing protein OS=Caenorhabditis elegans OX=6239 GN=rnf-1 PE=1 SV=2 |
| 9 | Q9C040 | 1.75e-04 | 42.7 | 744 | 0.154 | 0.032 | TRIM2_HUMAN Tripartite motif-containing protein 2 OS=Homo sapiens OX=9606 GN=TRIM2 PE=1 SV=1 |
| 10 | Q9ESN6 | 1.89e-04 | 42.7 | 744 | 0.154 | 0.032 | TRIM2_MOUSE Tripartite motif-containing protein 2 OS=Mus musculus OX=10090 GN=Trim2 PE=1 SV=1 |
| 11 | F1QFW2 | 2.04e-04 | 42.7 | 744 | 0.160 | 0.034 | F1QFW2_DANRE RING-type E3 ubiquitin transferase OS=Danio rerio OX=7955 GN=trim2 PE=3 SV=1 |
| 12 | D3ZQG6 | 2.16e-04 | 42.7 | 744 | 0.154 | 0.032 | TRIM2_RAT Tripartite motif-containing protein 2 OS=Rattus norvegicus OX=10116 GN=Trim2 PE=1 SV=2 |
| 13 | Q80VI1 | 3.95e-04 | 42.0 | 734 | 0.147 | 0.031 | TRI56_MOUSE E3 ubiquitin-protein ligase TRIM56 OS=Mus musculus OX=10090 GN=Trim56 PE=1 SV=1 |
| 14 | Q5PP23 | 0.001 | 40.8 | 562 | 0.141 | 0.039 | FLY1_ARATH Transmembrane E3 ubiquitin-protein ligase FLY1 OS=Arabidopsis thaliana OX=3702 GN=FLY1 PE=1 SV=1 |
| 15 | Q8IUD6 | 0.001 | 40.4 | 432 | 0.192 | 0.069 | RN135_HUMAN E3 ubiquitin-protein ligase RNF135 OS=Homo sapiens OX=9606 GN=RNF135 PE=1 SV=2 |
| 16 | Q9NXI6 | 0.001 | 39.7 | 227 | 0.154 | 0.106 | RN186_HUMAN E3 ubiquitin-protein ligase RNF186 OS=Homo sapiens OX=9606 GN=RNF186 PE=1 SV=1 |
| 17 | O17099 | 0.002 | 39.3 | 374 | 0.179 | 0.075 | O17099_CAEEL Major sperm protein OS=Caenorhabditis elegans OX=6239 GN=CELE_F42G2.5 PE=1 SV=1 |
| 18 | Q8N6D2 | 0.003 | 38.9 | 247 | 0.160 | 0.101 | RN182_HUMAN E3 ubiquitin-protein ligase RNF182 OS=Homo sapiens OX=9606 GN=RNF182 PE=1 SV=1 |
| 19 | Q9FY48 | 0.004 | 39.3 | 1625 | 0.147 | 0.014 | KEG_ARATH E3 ubiquitin-protein ligase KEG OS=Arabidopsis thaliana OX=3702 GN=KEG PE=1 SV=2 |
| 20 | P36406 | 0.006 | 38.5 | 574 | 0.199 | 0.054 | TRI23_HUMAN E3 ubiquitin-protein ligase TRIM23 OS=Homo sapiens OX=9606 GN=TRIM23 PE=1 SV=1 |

(a) Ranking of proteins is based on the E-value of a BLASTp alignment.
(b) ID1 and ID2 are sequence identities normalized by the query and template sequence lengths, respectively.

**Figure S5.** Structure and sequence templates used for StarFunc prediction of P0DH78.

#### Proteins with similar structure

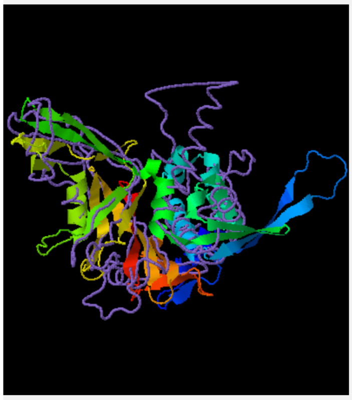

☐Spin On/Off

##### Top structural analogs (as identified by Foldseek and TM-align)

| Click to view | Rank | Structure template | TM-score1 | TM-score2 | Foldseek E-value | Length | ID1 | ID2 | Download | Description |
|---|---|---|---|---|---|---|---|---|---|---|
| ◉ | 1 | PDB:5cvnB | 0.450 | 0.438 | 3.040E-10 | 321 | 0.139 | 0.134 | 5cvnB.pdb.gz | Structural Insights into WD-Repeat 48 Activation of Ubiquitin-Specific Protease 46. |
| ○ | 2 | AFDB:Q09738 | 0.497 | 0.363 | 3.661E-10 | 449 | 0.155 | 0.107 | Q09738.pdb.gz | UBP8_SCHPO Probable ubiquitin carboxyl-terminal hydrolase 8 OS=Schizosaccharomyces pombe (strain 972 / ATCC 24843) OX=284812 GN=ubp8 PE=3 SV=1 |
| ○ | 3 | PDB:8a9kD | 0.457 | 0.490 | 4.615E-10 | 285 | 0.119 | 0.130 | 8a9kD.pdb.gz | Cryo-EM reveals a mechanism of USP1 inhibition through a cryptic binding site. |
| ○ | 4 | AFDB:B1AQJ2 | 0.432 | 0.148 | 1.071E-9 | 1098 | 0.155 | 0.044 | B1AQJ2.pdb.gz | UBP36_MOUSE Ubiquitin carboxyl-terminal hydrolase 36 OS=Mus musculus OX=10090 GN=Usp36 PE=1 SV=1 |
| ○ | 5 | AFDB:E7F4C0 | 0.443 | 0.150 | 1.833E-9 | 1104 | 0.145 | 0.041 | E7F4C0.pdb.gz | E7F4C0_DANRE Ubiquitin carboxyl-terminal hydrolase 36 OS=Danio rerio OX=7955 GN=usp37 PE=4 SV=2 |
| ○ | 6 | AFDB:P34547 | 0.454 | 0.352 | 1.833E-9 | 426 | 0.161 | 0.117 | P34547.pdb.gz | UBP46_CAEEL Ubiquitin carboxyl-terminal hydrolase 46 OS=Caenorhabditis elegans OX=6239 GN=usp-46 PE=1 SV=3 |
| ○ | 7 | AFDB:E7EYZ1 | 0.436 | 0.408 | 1.945E-9 | 337 | 0.100 | 0.092 | E7EYZ1.pdb.gz | E7EYZ1_DANRE Ubiquitin-specific peptidase 18 OS=Danio rerio OX=7955 GN=usp18 PE=4 SV=1 |
| ○ | 8 | AFDB:Q09879 | 0.435 | 0.150 | 2.470E-9 | 1108 | 0.135 | 0.038 | Q09879.pdb.gz | UBP72_SCHPO Ubiquitin carboxyl-terminal hydrolase 5 OS=Schizosaccharomyces pombe (strain 972 / ATCC 24843) OX=284812 GN=ubp5 PE=1 SV=3 |
| ○ | 9 | AFDB:Q583H3 | 0.489 | 0.350 | 3.328E-9 | 465 | 0.126 | 0.084 | Q583H3.pdb.gz | Q583H3_TRYB2 Ubiquitin carboxyl-terminal hydrolase OS=Trypanosoma brucei brucei (strain 927/4 GUTat10.1) OX=185431 GN=Tb04.30O21.180 PE=3 SV=1 |
| ○ | 10 | AFDB:Q9SB51 | 0.468 | 0.171 | 3.532E-9 | 1008 | 0.110 | 0.034 | Q9SB51.pdb.gz | UBP16_ARATH Ubiquitin carboxyl-terminal hydrolase 16 OS=Arabidopsis thaliana OX=3702 GN=UBP16 PE=1 SV=1 |
| ○ | 11 | AFDB:Q9WTV6 | 0.396 | 0.345 | 4.484E-9 | 368 | 0.152 | 0.128 | Q9WTV6.pdb.gz | UBP18_MOUSE Ubl carboxyl-terminal hydrolase 18 OS=Mus musculus OX=10090 GN=Usp18 PE=1 SV=2 |
| ○ | 12 | AFDB:Q8MQX4 | 0.422 | 0.176 | 4.760E-9 | 896 | 0.116 | 0.040 | Q8MQX4.pdb.gz | Q8MQX4_DROME Ubiquitin carboxyl-terminal hydrolase OS=Drosophila melanogaster OX=7227 GN=Usp8 PE=2 SV=1 |
| ○ | 13 | PDB:4fjcE | 0.414 | 0.312 | 7.176E-9 | 434 | 0.148 | 0.106 | 4fjcE.pdb.gz | A Role for Intersubunit Interactions in Maintaining SAGA Deubiquitinating Module Structure and Activity. |
| ○ | 14 | PDB:5k1bA | 0.454 | 0.514 | 1.089E-8 | 264 | 0.126 | 0.148 | 5k1bA.pdb.gz | Allosteric Activation of Ubiquitin-Specific Proteases by beta-Propeller Proteins UAF1 and WDR20. |
| ○ | 15 | PDB:3mhsA | 0.504 | 0.364 | 2.666E-8 | 455 | 0.148 | 0.101 | 3mhsA.pdb.gz | Structural insights into the assembly and function of the SAGA deubiquitinating module. |
| ○ | 16 | PDB:7yxxA | 0.450 | 0.103 | 3.188E-8 | 1754 | 0.177 | 0.031 | 7yxxA.pdb.gz | |
| ○ | 17 | PDB:7ay1D | 0.433 | 0.400 | 3.384E-8 | 341 | 0.132 | 0.120 | 7ay1D.pdb.gz | Structural basis of FANCD2 deubiquitination by USP1-UAF1. |
| ○ | 18 | PDB:5k16B | 0.422 | 0.402 | 3.813E-8 | 328 | 0.139 | 0.131 | 5k16B.pdb.gz | Allosteric Activation of Ubiquitin-Specific Proteases by beta-Propeller Proteins UAF1 and WDR20. |
| ○ | 19 | PDB:5gjqx | 0.453 | 0.407 | 6.522E-8 | 355 | 0.113 | 0.099 | 5gjqx.pdb.gz | An atomic structure of the human 26S proteasome. |
| ○ | 20 | PDB:7qo48 | 0.493 | 0.424 | 1.184E-7 | 372 | 0.142 | 0.118 | 7qo48.pdb.gz | Allosteric control of Ubp6 and the proteasome via a bidirectional switch. |

(a) The query structure is shown in cartoon, while the structural analog is displayed using a backbone trace.
(b) Ranking of proteins is based on E-value of Foldseek alignment.
(c) TM-score1 and ID1 are TM-score and sequence identity normalized by query sequence length.
(d) TM-score2 and ID2 are TM-score and sequence identity normalized by template sequence length.

#### Proteins with similar sequence

##### Top sequence homologs in UniProt

| Rank | Template | E-value | Bit-score | Length | ID1 | ID2 | Description |
|---|---|---|---|---|---|---|---|
| 1 | H2KYE0 | 0.13 | 36.6 | 1123 | 0.071 | 0.020 | NRA4_CAEEL Nicotinic receptor-associated protein 4 OS=Caenorhabditis elegans OX=6239 GN=nra-4 PE=1 SV=1 |
| 2 | Q9QYE2 | 0.78 | 33.9 | 670 | 0.065 | 0.030 | SO1A6_RAT Solute carrier organic anion transporter family member 1A6 OS=Rattus norvegicus OX=10116 GN=Slco1a6 PE=2 SV=1 |
| 3 | Q9EP96 | 3.2 | 32.0 | 670 | 0.065 | 0.030 | SO1A4_MOUSE Solute carrier organic anion transporter family member 1A4 OS=Mus musculus OX=10090 GN=Slco1a4 PE=1 SV=1 |
| 4 | Q9QXZ6 | 4.0 | 31.6 | 670 | 0.068 | 0.031 | SO1A1_MOUSE Solute carrier organic anion transporter family member 1A1 OS=Mus musculus OX=10090 GN=Slco1a1 PE=1 SV=1 |
| 5 | P53065 | 5.3 | 31.2 | 400 | 0.052 | 0.040 | TAD1_YEAST tRNA-specific adenosine deaminase 1 OS=Saccharomyces cerevisiae (strain ATCC 204508 / S288c) OX=559292 GN=TAD1 PE=1 SV=1 |
| 6 | P46720 | 6.7 | 30.8 | 670 | 0.061 | 0.028 | SO1A1_RAT Solute carrier organic anion transporter family member 1A1 OS=Rattus norvegicus OX=10116 GN=Slco1a1 PE=1 SV=1 |
| 7 | O65581 | 7.1 | 30.8 | 358 | 0.110 | 0.095 | ALFC5_ARATH Fructose-bisphosphate aldolase 5, cytosolic OS=Arabidopsis thaliana OX=3702 GN=FBA5 PE=1 SV=1 |
| 8 | P04925 | 9.2 | 30.0 | 254 | 0.035 | 0.043 | PRIO_MOUSE Major prion protein OS=Mus musculus OX=10090 GN=Prnp PE=1 SV=2 |

(a) Ranking of proteins is based on the E-value of a BLASTp alignment.
(b) ID1 and ID2 are sequence identities normalized by the query and template sequence lengths, respectively.

**Figure S6.** Structure and sequence templates used for StarFunc prediction of Q4W4Y0.

#### Proteins with similar structure

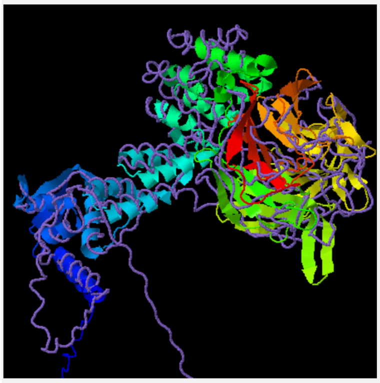

Spin On/Off

##### Top structural analogs (as identified by Foldseek and TM-align)

| Click to view | Rank | Structure template | TM-score1 | TM-score2 | Foldseek E-value | Length | ID1 | ID2 | Download | Description |
|---|---|---|---|---|---|---|---|---|---|---|
| | 1 | AFDB:Q9Y2M5 | 0.736 | 0.697 | 1.378E-70 | 609 | 0.427 | 0.401 | Q9Y2M5.pdb.gz | KLH20_HUMAN Kelch-like protein 20 OS=Homo sapiens OX=9606 GN=KLHL20 PE=1 SV=4 |
| | 2 | AFDB:Q6TDP4 | 0.535 | 0.480 | 6.173E-70 | 642 | 0.412 | 0.366 | Q6TDP4.pdb.gz | KLH17_HUMAN Kelch-like protein 17 OS=Homo sapiens OX=9606 GN=KLHL17 PE=1 SV=1 |
| | 3 | AFDB:Q9VUU5 | 0.579 | 0.539 | 6.173E-70 | 623 | 0.424 | 0.388 | Q9VUU5.pdb.gz | KLHDB_DROME Kelch-like protein diablo OS=Drosophila melanogaster OX=7227 GN=dbo PE=1 SV=1 |
| | 4 | AFDB:Q9JI74 | 0.569 | 0.449 | 1.620E-67 | 751 | 0.398 | 0.302 | Q9JI74.pdb.gz | KLHL1_MOUSE Kelch-like protein 1 OS=Mus musculus OX=10090 GN=Klhl1 PE=2 SV=2 |
| | 5 | AFDB:E9Q4F2 | 0.528 | 0.526 | 2.623E-67 | 574 | 0.373 | 0.371 | E9Q4F2.pdb.gz | KLH18_MOUSE Kelch-like protein 18 OS=Mus musculus OX=10090 GN=Klhl18 PE=1 SV=1 |
| | 6 | AFDB:A5PKX1 | 0.627 | 0.518 | 8.076E-67 | 718 | 0.380 | 0.302 | A5PKX1.pdb.gz | A5PKX1_HUMAN Kelch-like 4 (Drosophila) OS=Homo sapiens OX=9606 GN=KLHL4 PE=2 SV=1 |
| | 7 | AFDB:Q53G59 | 0.827 | 0.831 | 2.118E-66 | 568 | 0.385 | 0.387 | Q53G59.pdb.gz | KLH12_HUMAN Kelch-like protein 12 OS=Homo sapiens OX=9606 GN=KLHL12 PE=1 SV=2 |
| | 8 | AFDB:O95198 | 0.498 | 0.481 | 3.817E-66 | 593 | 0.371 | 0.358 | O95198.pdb.gz | KLHL2_HUMAN Kelch-like protein 2 OS=Homo sapiens OX=9606 GN=KLHL2 PE=1 SV=2 |
| | 9 | AFDB:Q96PQ7 | 0.593 | 0.467 | 1.308E-65 | 755 | 0.389 | 0.294 | Q96PQ7.pdb.gz | KLHL5_HUMAN Kelch-like protein 5 OS=Homo sapiens OX=9606 GN=KLHL5 PE=1 SV=3 |
| | 10 | AFDB:E0CZ16 | 0.546 | 0.534 | 1.903E-65 | 587 | 0.373 | 0.363 | E0CZ16.pdb.gz | KLHL3_MOUSE Kelch-like protein 3 OS=Mus musculus OX=10090 GN=Klhl3 PE=1 SV=2 |
| | 11 | PDB:2uvkB | 0.408 | 0.648 | 8.885E-17 | 349 | 0.095 | 0.155 | 2uvkB.pdb.gz | |
| | 12 | PDB:5gqtA | 0.369 | 0.445 | 1.161E-16 | 470 | 0.109 | 0.132 | 5gqtA.pdb.gz | |
| | 13 | PDB:5gq0A | 0.376 | 0.613 | 8.426E-15 | 338 | 0.098 | 0.166 | 5gq0A.pdb.gz | |
| | 14 | PDB:2zwaB | 0.368 | 0.312 | 1.519E-14 | 683 | 0.093 | 0.078 | 2zwaB.pdb.gz | Structural basis of tRNA modification with CO2 fixation and methylation by wybutosine synthesizing enzyme TYW4. |
| | 15 | PDB:6wcqC | 0.195 | 0.813 | 4.684E-9 | 130 | 0.079 | 0.346 | 6wcqC.pdb.gz | |
| | 16 | PDB:2hqsA | 0.365 | 0.486 | 9.396E-9 | 412 | 0.040 | 0.056 | 2hqsA.pdb.gz | |
| | 17 | PDB:3jcmB | 0.302 | 0.386 | 3.782E-8 | 429 | 0.109 | 0.145 | 3jcmB.pdb.gz | The 3.8 angstrom structure of the U4/U6.U5 tri-snRNP: Insights into spliceosome assembly and catalysis |
| | 18 | PDB:4v7fq | 0.271 | 0.332 | 5.215E-8 | 443 | 0.060 | 0.077 | 4v7fq.pdb.gz | 60S ribosome biogenesis requires rotation of the 5S ribonucleoprotein particle. |
| | 19 | PDB:7z8bF | 0.252 | 0.303 | 5.215E-8 | 457 | 0.074 | 0.092 | 7z8bF.pdb.gz | Structure of CRL7 FBXW8 reveals coupling with CUL1-RBX1/ROC1 for multi-cullin-RING E3-catalyzed ubiquitin ligation. |
| | 20 | PDB:7mq8SG | 0.316 | 0.435 | 8.909E-8 | 389 | 0.072 | 0.105 | 7mq8SG.pdb.gz | Nucleolar maturation of the human small subunit processome. |

(a) The query structure is shown in cartoon, while the structural analog is displayed using a backbone trace.
(b) Ranking of proteins is based on E-value of Foldseek alignment.
(c) TM-score1 and ID1 are TM-score and sequence identity normalized by query sequence length.
(d) TM-score2 and ID2 are TM-score and sequence identity normalized by template sequence length.

#### Proteins with similar sequence

##### Top sequence homologs in UniProt

| Rank | Template | E-value | Bit-score | Length | ID1 | ID2 | Description |
|---|---|---|---|---|---|---|---|
| 1 | Q9Y2M5 | 5.97e-156 | 461 | 609 | 0.426 | 0.399 | KLH20_HUMAN Kelch-like protein 20 OS=Homo sapiens OX=9606 GN=KLHL20 PE=1 SV=4 |
| 2 | Q9VUU5 | 2.38e-148 | 442 | 623 | 0.419 | 0.384 | KLHDB_DROME Kelch-like protein diablo OS=Drosophila melanogaster OX=7227 GN=dbo PE=1 SV=1 |
| 3 | Q8K430 | 2.48e-145 | 435 | 640 | 0.417 | 0.372 | KLH17_RAT Kelch-like protein 17 OS=Rattus norvegicus OX=10116 GN=Klhl17 PE=1 SV=1 |
| 4 | Q6TDP4 | 7.52e-145 | 434 | 642 | 0.415 | 0.369 | KLH17_HUMAN Kelch-like protein 17 OS=Homo sapiens OX=9606 GN=KLHL17 PE=1 SV=1 |
| 5 | Q96PQ7 | 7.63e-143 | 432 | 755 | 0.391 | 0.295 | KLHL5_HUMAN Kelch-like protein 5 OS=Homo sapiens OX=9606 GN=KLHL5 PE=1 SV=3 |
| 6 | Q9JI74 | 3.38e-134 | 410 | 751 | 0.396 | 0.301 | KLHL1_MOUSE Kelch-like protein 1 OS=Mus musculus OX=10090 GN=Klhl1 PE=2 SV=2 |
| 7 | Q9NR64 | 3.65e-134 | 410 | 748 | 0.398 | 0.303 | KLHL1_HUMAN Kelch-like protein 1 OS=Homo sapiens OX=9606 GN=KLHL1 PE=1 SV=1 |
| 8 | Q5U374 | 5.98e-134 | 404 | 564 | 0.380 | 0.385 | KLH12_DANRE Kelch-like protein 12 OS=Danio rerio OX=7955 GN=klhl12 PE=2 SV=2 |
| 9 | E9Q4F2 | 1.04e-132 | 400 | 574 | 0.373 | 0.371 | KLH18_MOUSE Kelch-like protein 18 OS=Mus musculus OX=10090 GN=Klhl18 PE=1 SV=1 |
| 10 | O94889 | 3.09e-132 | 399 | 574 | 0.371 | 0.369 | KLH18_HUMAN Kelch-like protein 18 OS=Homo sapiens OX=9606 GN=KLHL18 PE=1 SV=3 |
| 11 | Q9UH77 | 5.27e-132 | 399 | 587 | 0.378 | 0.368 | KLHL3_HUMAN Kelch-like protein 3 OS=Homo sapiens OX=9606 GN=KLHL3 PE=1 SV=2 |
| 12 | E0CZ16 | 3.27e-130 | 395 | 587 | 0.375 | 0.365 | KLHL3_MOUSE Kelch-like protein 3 OS=Mus musculus OX=10090 GN=Klhl3 PE=1 SV=2 |
| 13 | Q53G59 | 1.72e-129 | 392 | 568 | 0.382 | 0.384 | KLH12_HUMAN Kelch-like protein 12 OS=Homo sapiens OX=9606 GN=KLHL12 PE=1 SV=2 |
| 14 | A5PKX1 | 6.02e-126 | 388 | 718 | 0.387 | 0.308 | A5PKX1_HUMAN Kelch-like 4 (Drosophila) OS=Homo sapiens OX=9606 GN=KLHL4 PE=2 SV=1 |
| 15 | Q9C0H6 | 6.02e-126 | 388 | 718 | 0.387 | 0.308 | KLHL4_HUMAN Kelch-like protein 4 OS=Homo sapiens OX=9606 GN=KLHL4 PE=1 SV=2 |
| 16 | Q8JZP3 | 1.67e-125 | 383 | 593 | 0.375 | 0.361 | KLHL2_MOUSE Kelch-like protein 2 OS=Mus musculus OX=10090 GN=Klhl2 PE=2 SV=1 |
| 17 | O95198 | 1.71e-125 | 383 | 593 | 0.373 | 0.359 | KLHL2_HUMAN Kelch-like protein 2 OS=Homo sapiens OX=9606 GN=KLHL2 PE=1 SV=2 |
| 18 | Q04652 | 9.14e-125 | 401 | 1477 | 0.373 | 0.144 | KELC_DROME Ring canal kelch protein OS=Drosophila melanogaster OX=7227 GN=kel PE=1 SV=4 |
| 19 | Q9VGE5 | 1.27e-118 | 364 | 575 | 0.359 | 0.357 | Q9VGE5_DROME Kelch-like protein diablo OS=Drosophila melanogaster OX=7227 GN=KLHL18 PE=2 SV=1 |
| 20 | Q9Z2X8 | 5.44e-116 | 359 | 624 | 0.359 | 0.329 | KEAP1_MOUSE Kelch-like ECH-associated protein 1 OS=Mus musculus OX=10090 GN=Keap1 PE=1 SV=1 |

(a) Ranking of proteins is based on the E-value of a BLASTp alignment.
(b) ID1 and ID2 are sequence identities normalized by the query and template sequence lengths, respectively.

**Figure S7.** Structure and sequence templates used for StarFunc prediction of Q9NXS3.

#### Supporting Tables

**Table S1.** Fmax, Smin, and wFmax values for StarFunc and existing comparison methods applied to the test set. Values that are the best in each category are highlighted by bold fonts.

| Method | Fmax | | | | Smin | | | | wFmax | | | |
|---|---|---|---|---|---|---|---|---|---|---|---|---|
| | MF | BP | CC | All | MF | BP | CC | All | MF | BP | CC | All |
| StarFunc | **0.647** | **0.465** | **0.819** | **0.643** | **5.848** | **20.16** | **4.034** | **10.02** | **0.639** | **0.445** | **0.729** | **0.604** |
| hfm7zc * | 0.634 | 0.452 | 0.791 | 0.625 | 6.077 | 20.712 | 4.300 | 10.363 | 0.616 | 0.433 | 0.698 | 0.582 |
| SPROF-GO | 0.596 | 0.398 | 0.773 | 0.589 | 6.421 | 20.80 | 4.467 | 10.56 | 0.570 | 0.375 | 0.670 | 0.539 |
| ATGO+ | 0.453 | 0.430 | 0.762 | 0.548 | 6.220 | 20.21 | 5.136 | 10.52 | 0.400 | 0.411 | 0.661 | 0.490 |
| TALE+ | 0.495 | 0.339 | 0.731 | 0.522 | 7.072 | 22.09 | 5.308 | 11.49 | 0.472 | 0.317 | 0.608 | 0.466 |
| GoFDR | 0.505 | 0.369 | 0.632 | 0.502 | 7.963 | 24.01 | 7.560 | 13.18 | 0.476 | 0.353 | 0.478 | 0.435 |
| DeepGOplus | 0.489 | 0.312 | 0.709 | 0.503 | 7.580 | 23.12 | 6.361 | 12.35 | 0.438 | 0.309 | 0.551 | 0.433 |
| AnnoPRO | 0.435 | 0.353 | 0.677 | 0.488 | 8.054 | 22.38 | 6.333 | 12.26 | 0.396 | 0.337 | 0.540 | 0.424 |
| ProteInfer | 0.397 | 0.359 | 0.605 | 0.454 | 8.535 | 23.38 | 7.304 | 13.04 | 0.376 | 0.339 | 0.454 | 0.390 |
| COFACTOR | 0.363 | 0.369 | 0.668 | 0.467 | 6.804 | 21.53 | 6.562 | 11.63 | 0.340 | 0.350 | 0.508 | 0.401 |
| GOtcha | 0.408 | 0.267 | 0.648 | 0.441 | 7.322 | 22.87 | 6.693 | 12.30 | 0.387 | 0.246 | 0.458 | 0.364 |
| DeepGO-SE | 0.561 | 0.325 | 0.325 | 0.404 | 6.810 | 22.21 | 8.192 | 12.40 | 0.535 | 0.301 | 0.127 | 0.321 |
| DeepFRI | 0.274 | 0.239 | 0.562 | 0.358 | 8.071 | 23.37 | 6.945 | 12.80 | 0.243 | 0.224 | 0.402 | 0.290 |

* "hfm7zc" refers to the preliminary implementation of StarFunc used by CAFA5, while "StarFunc" refers to the final version of StarFunc.

**Table S2.** Fmax, Smin and wFmax values for StarFunc and its component methods on the validation set. Values that are the best in each category are highlighted by bold fonts.

| Method | Fmax | | | | Smin | | | | wFmax | | | |
|---|---|---|---|---|---|---|---|---|---|---|---|---|
| | MF | BP | CC | All | MF | BP | CC | All | MF | BP | CC | All |
| StarFunc | **0.625** | **0.434** | **0.694** | **0.584** | 6.304 | **18.27** | **5.854** | **10.14** | **0.609** | **0.411** | **0.608** | **0.542** |
| InterLabelGO | 0.618 | 0.421 | 0.685 | 0.575 | 6.550 | 18.58 | 6.106 | 10.41 | 0.596 | 0.399 | 0.594 | 0.530 |
| Sequence | 0.583 | 0.372 | 0.642 | 0.532 | 6.876 | 20.54 | 6.868 | 11.43 | 0.560 | 0.350 | 0.535 | 0.481 |
| Structure | 0.598 | 0.336 | 0.625 | 0.520 | 6.870 | 21.76 | 6.972 | 11.87 | 0.572 | 0.316 | 0.509 | 0.466 |
| Pfam | 0.529 | 0.311 | 0.575 | 0.472 | 7.374 | 22.30 | 7.465 | 12.38 | 0.512 | 0.289 | 0.471 | 0.424 |
| PPI | 0.416 | 0.330 | 0.588 | 0.444 | 9.577 | 21.44 | 7.226 | 12.75 | 0.368 | 0.300 | 0.497 | 0.388 |
| StarFunc (no InterLabelGO) | 0.597 | 0.391 | 0.655 | 0.548 | 6.614 | 20.15 | 6.588 | 11.12 | 0.578 | 0.370 | 0.561 | 0.503 |
| StarFunc (no sequence) | 0.597 | 0.391 | 0.655 | 0.548 | 6.614 | 20.15 | 6.588 | 11.12 | 0.578 | 0.370 | 0.561 | 0.503 |
| StarFunc (no structure) | 0.620 | 0.431 | 0.694 | 0.582 | 6.443 | 18.34 | 5.870 | 10.22 | 0.603 | 0.407 | **0.608** | 0.540 |
| StarFunc (no Pfam) | **0.625** | 0.433 | 0.693 | **0.584** | 6.332 | 18.28 | 5.870 | 10.16 | 0.608 | 0.410 | **0.608** | **0.542** |
| StarFunc (no PPI) | 0.622 | 0.419 | 0.682 | 0.575 | **6.290** | 18.74 | 5.952 | 10.32 | 0.608 | 0.398 | 0.595 | 0.533 |

**Table S3.** Summary of PRAME family proteins among missing proteins that are predicted by StarFunc to be associated with the Cul2-RING ubiquitin ligase complex.

| UniProt accession | Protein Name | Gene Name | Prediction score for GO:0031462 "Cul2-RING ubiquitin ligase complex" |
|---|---|---|---|
| Q5TYX0 | PRAME family member 5 | PRAMEF5 | 0.994 |
| O60813 | PRAME family member 11 | PRAMEF11 | 0.992 |
| A6NGN4 | PRAME family member 25 | PRAMEF25 | 0.991 |
| O60810 | PRAME family member 4 | PRAMEF4 | 0.991 |
| A3QJZ7 | PRAME family member 27 | PRAMEF27 | 0.988 |
| Q5VT98 | PRAME family member 20 | PRAMEF20 | 0.988 |
| Q5VWM4 | PRAME family member 8 | PRAMEF8 | 0.822 |
| O95522 | PRAME family member 12 | PRAMEF12 | 0.810 |
| Q5VXH5 | PRAME family member 7 | PRAMEF7 | 0.808 |
| A3QJZ6 | PRAME family member 22 | PRAMEF22 | 0.737 |
| Q5VWM3 | PRAME family member 18 | PRAMEF18 | 0.728 |
| Q5SWL8 | PRAME family member 19 | PRAMEF19 | 0.727 |
